# Founders predict trait evolution and population performance after evolutionary rescue in the red flour beetle

**DOI:** 10.1101/2025.03.17.640637

**Authors:** Vrinda Ravi Kumar, Shyamsunder Buddh, Shivansh Singhal, Arun Prakash, Deepa Agashe

**Affiliations:** National Centre for Biological Sciences, Tata Institute of Fundamental Research, Bellary Road, Bangalore, India 560065

**Keywords:** Evolutionary rescue, adaptation, *Tribolium castaneum*, red flour beetle, phenotypic variation, rapid evolution

## Abstract

Evolutionary rescue helps populations survive environmental change, but the phenotypic and demographic factors associated with rescue dynamics and its long-term effects remain unclear. We experimentally evolved 10 wild-collected populations of flour beetles from across India in a suboptimal corn resource for 70 generations (>5 years), collecting >10,000 population census points book-ended by measurements of fitness-related traits for 30 experimental lines. Despite clear ancestral trait differences, all lines showed highly parallel evolutionary rescue within 20 generations. Long-term average population size varied across source populations and was positively correlated with ancestral development rate, which increased convergently across populations and emerged as the single best predictor of population performance during and after evolutionary rescue. Notably, specific demographic events during rescue (such as the rate of population decline and recovery) were uncorrelated both with ancestral trait distributions and post-rescue adaptation. Our results support prior work showing founder traits as key predictors of adaptation, and highlight their role in long-term adaptation and trait evolution following evolutionary rescue.

**SIGNIFICANCE STATEMENT:** Understanding how populations adapt to sudden environmental change is vital given accelerating biodiversity declines, the rise of antimicrobial resistance and climate change. We chart evolutionary rescue — where rapid adaptation prevents extinction — in flour beetles across 70 generations. Using >10,000 census points and measuring fitness in ancestors, their offspring and in evolved lines, we show that larval development rate is the strongest correlate of long-term population success across diverse wild-collected populations. The effect of ancestral variation is dynamic and disassociated from finer-grained details of the rescue. All populations show highly parallel evolutionary rescue, with convergent evolution in key traits. These findings have direct implications for predicting species’ survival after environmental change, ranging from conservation to pest and pathogen management.

## INTRODUCTION

Abrupt environmental changes (e.g., climate change or urbanization) can cause species abundance to decline rapidly, potentially causing extinction (1, 2) and reduction in local biodiversity (3, 4). Evolutionary rescue — likely an important mechanism driving species persistence in such conditions (1, 5, 6) — refers to the demographic recovery of declining populations due to rapid adaptive evolution, with a hallmark U-shaped curve of population size over time (7–10). Evolutionary rescue is aided by several genetic and demographic factors (7, 11–16), such as founding population size (17–21), standing genetic variation (20, 22–24), and demographic history (25–28). Although rescue has wide-ranging applications in conservation, pest control and medicine (7), detecting and predicting evolutionary rescue outside the laboratory is fraught with logistical and technical challenges (29). Thus, despite the large body of work on this topic, some important gaps remain in our empirical understanding of evolutionary rescue.

First, evolutionary rescue is often framed as a binary phenomenon: populations either go extinct or are rescued. Thus, most studies focus on factors that influence the probability of successful rescue, when mean absolute fitness (which initially declines in the new environment) increases beyond 1 via adaptive evolution (7, 8, 10, 12, 29, 30). However, the detailed demographic changes during rescue, and longer-term consequences of the rescue process, have received relatively little attention (but see (10, 28, 31)). For instance, the frequency of rare adaptive alleles and the waiting time to sample new beneficial mutations should influence the bottleneck period after the initial population crash, the slope of the population size rebound, and how much variation is retained (7, 8, 10, 12, 29). These processes should depend on the mean and variance in founder traits (which determine the response to selection during rescue), and the outcomes may in turn influence population performance after rescue. But empirically, we do not know what demographic factors and traits determine population size during and after successful rescue, and whether the speed or the magnitude of adaptation during successful rescue predicts subsequent population dynamics and adaptation. This knowledge is critical for effective applications of rescue, particularly in conservation. However, because most studies of evolutionary rescue focus on short-term binary outcomes, these questions remain unanswered.

Second, the phenotypic drivers of evolutionary rescue and subsequent population performance are not always understood or predictable, and yet are vitally important to understand rescue (32). The broadly beneficial role of founding genetic diversity in rescue (20, 22–24, 31, 33) is driven by genetic variation which contributes to phenotypic traits that increase survival and reproduction in the new environment (34, 35), and that are therefore directly visible to selection (36). Prior experiments suggest that even in populations with high initial diversity, only some founder traits may contribute to rescue and adult population size after recovery (31). But exactly which traits of founding individuals are relevant for the initial rate of population decline or the speed of evolutionary rescue, and how long do these effects persist?

Answering these questions is difficult for several reasons. Selection may simultaneously act on multiple phenotypic traits in the new environment, which may also interact with each other, e.g., via genetic correlations (37). Trait distributions can change during rescue as selection pressures change, or due to demographic and genetic stochasticity imposed by rapid initial population decline in the new environment (8, 12). For instance, high mortality in the new environment can immediately and strongly change trait distributions (38). Finally, trait distributions and population dynamics can be influenced by plasticity (“plastic rescue”) (32, 39–46) and carry-over (e.g., maternal) effects of founder individuals (47–50). For instance, short-term population dynamics in the red flour beetle *Tribolium castaneum* depend on both the past resource in which colonizing individuals developed, and the newly colonized resource (46). Therefore, identifying which founder phenotypes are critical for evolutionary rescue and later population performance requires the measurement of trait distributions immediately after introduction to the new habitat (which likely includes both single-generation selective mortality and the impact of plasticity), as well as after rescue (after long-term evolution). Very few studies examine multi-trait responses to abrupt environmental change (51), though such data are critical to predict both the probability and outcome of rescue.

We used laboratory experimental evolution in a harsh novel resource (corn flour), founding each line with one of ten wild-collected populations of the red flour beetle *Tribolium castaneum* with high fitness in their ancestral wheat flour environment (Fig. 1). Prior to starting the experiment, we first confirmed that these ten source populations had distinct distributions of several fitness-related traits, allowing the possibility of divergent outcomes during rescue and long-term adaptation. We then collected complete census data from each evolving line for 70 generation cycles (280 weeks, or 5 years and 4 months), propagated in an overlapping (rather than discrete) generation design. We observed successful evolutionary rescue in nearly all populations within 20 generations, allowing us to monitor the impacts of ancestral traits for about 50 generations after rescue. Using our census data, we tested which founder traits may have driven specific demographic events during evolutionary rescue (e.g., the decline, bottleneck and recovery phases) and which traits predicted long term population performance (population size and stability). We also collected phenotypic data to test whether and which trait distributions changed immediately in the new habitat (after source populations spent one generation in corn flour) and after relatively long-term evolution (70 generations). Although we use the term ‘founders’, our work is relevant not only in the context of colonization of new habitats, but also *in situ* environmental change. We identify specific fitness components with surprisingly far-reaching effects on long term population performance, with potential applications for predicting population fate in the wild (29).

**Figure 1.**
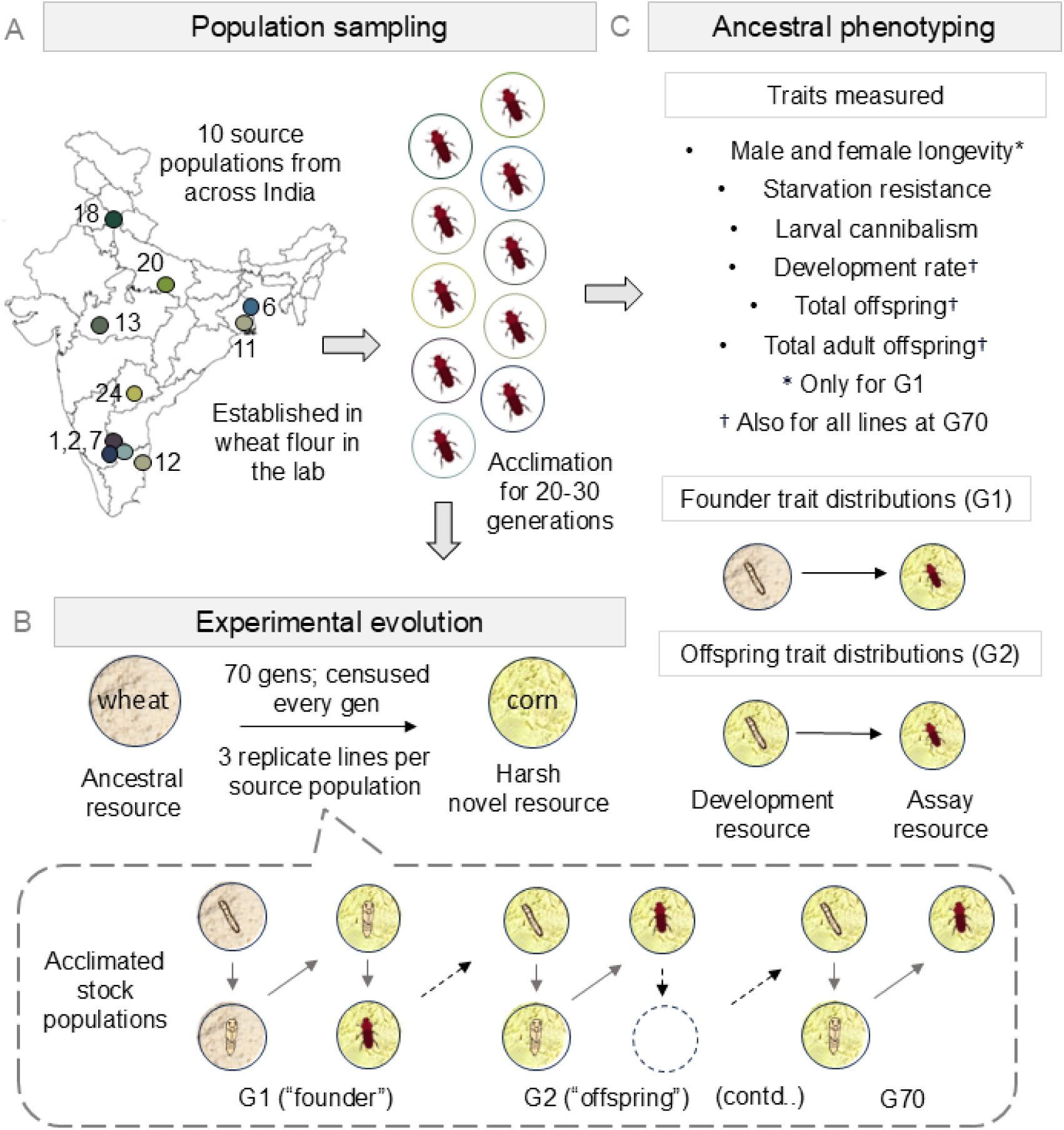
Schematic of the experimental design. (A) Population sampling and lab acclimation. Ten source populations were collected from across the Indian mainland (marked on the map: Survey of India), brought into the lab and acclimated to lab conditions in wheat flour (ancestral resource) for 20-30 generations. (B) Experimental evolution. We initiated selection lines in corn flour (a harsh novel environment), and transferred lines every 4 weeks (1 generation cycle) to fresh corn flour, for 70 generation cycles (280 weeks). (C) Ancestral phenotyping of the source populations. We measured seven fitness-related traits after rearing stock-derived individuals under two conditions – one mimicking the founders of the lines (generation 0, G0), and the other mimicking their immediate offspring that developed in corn (G1). We also measured three of these traits indicated by our results (see ‘Results’) in all evolved lines at generation cycle 70.

## RESULTS

### Source populations had distinct ancestral trait distributions

We first measured founder phenotypes for the ten source populations (Fig. S1; Supplementary Methods) collected from eight different locations around India (Fig. 1A). To capture the phenotypic trait distribution of the founders (individuals that developed in wheat and were then introduced to corn; Fig. 1B), we used wheat-developed individuals from the source populations exposed to corn for the first time, which mimicked the developmental environment of the actual founders used to initiate the selection lines (Fig. 1C). Note that we did not measure traits of the actual founder individuals since the phenotyping would interfere with the experimental evolution. We focused on several traits relevant for flour beetle fitness, broadly affecting survival (adult longevity/survivorship and starvation resistance), reproduction (total offspring, and total adult offspring) and other aspects of fitness that indirectly contribute to survival, reproduction, or both (larval cannibalism and offspring development rate).

Source populations were phenotypically distinct for all measured traits, except total adult offspring produced per female (Fig. 2; see Table S1 for full statistical report). The differences were substantial, with adults in some populations living for about 1.5 days longer than others under starvation, and females from some populations producing four times as many total offspring as others. In the slowest-developing populations, only approximately 3% of offspring pupated within the experimental window (4 weeks), compared to about 42% in the fastest-developing populations. Given these ecologically relevant differences across source population trait distributions, we expected to find divergent adaptive trajectories during rescue and subsequent trait evolution in corn. For instance, offspring development rate (hereafter ‘development rate’) and total offspring are important traits that directly affect population size. These traits were most strongly affected by source population identity (Table S1), suggesting that they could drive variation in population performance during the early stages of rescue and adaptation.

**Figure 2.**
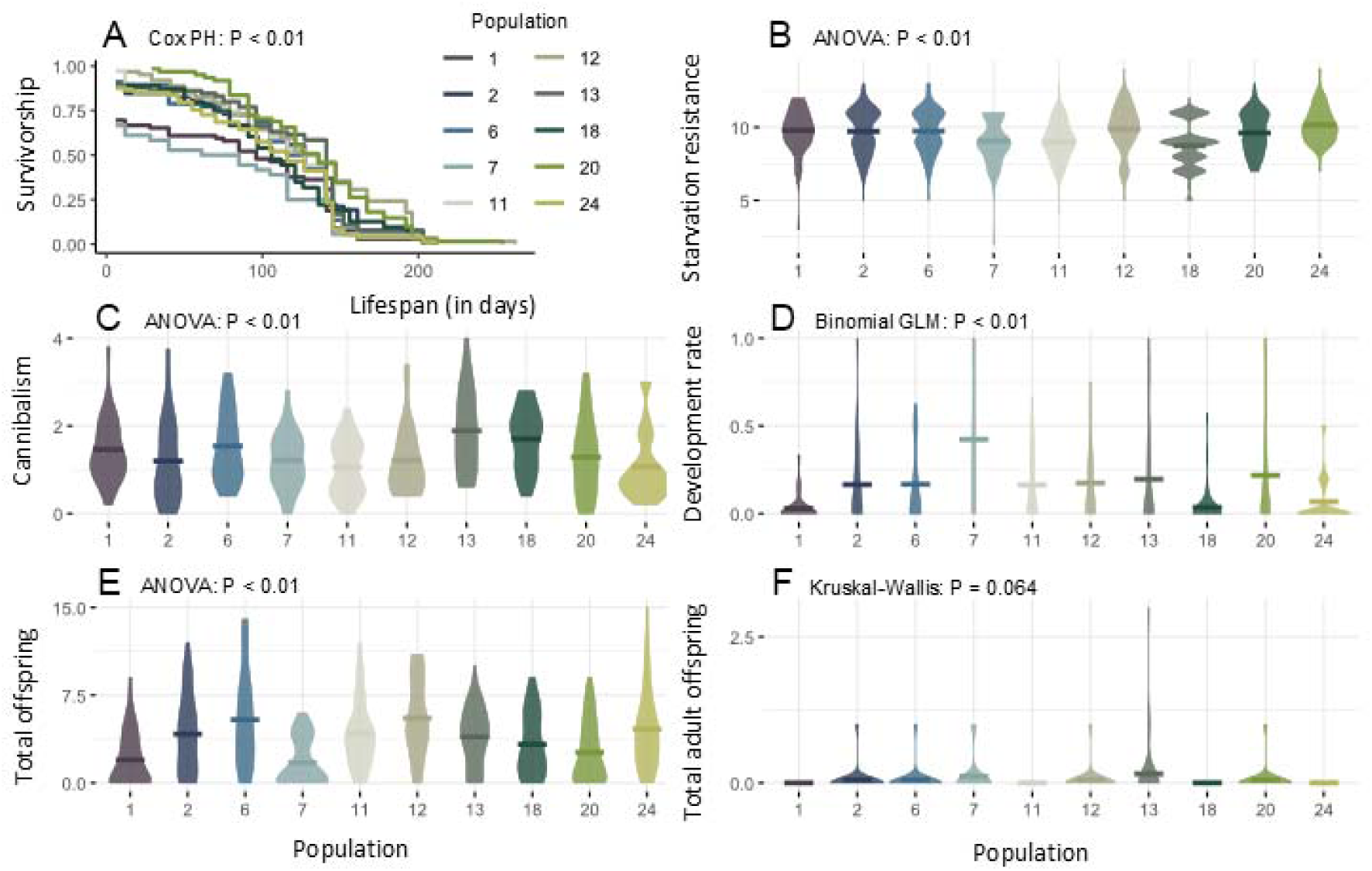
Variation in ancestral phenotypic traits across the ten source populations. Phenotypic trait distributions for each source population for different fitness-related traits (panels A-F). Solid lines inside the violin plots represent distribution means for different populations. (A) Beetle longevity (in days, males and females equally represented; N = 60-80 individuals/population) represented as Kaplan-Meier survivorship curves (B) Starvation resistance (in days, n = 48 pupae per population); (C) Per-capita larval cannibalism (eggs consumed/larvae, n = 25 per population), (D) Offspring development rate (proportion of pupated or eclosed individuals, n = 25-40 females per population) and (E–F) measures of reproductive fitness per female (n = 25-40 females per population), (E) Number of total offspring, and (F) Number of adult offspring. Founder starvation resistance assays for population 13 (panel B) were lost to an accident. One extreme data point is not shown in panel E (population 12, 42 total offspring) for ease of visualization, but is included in the reported statistics. We used different tests to assess whether source population had a significant impact on founder trait distributions – these varied by trait and are reported directly in the relevant panels. See Table S1 for summary statistics and Methods for trait assay details.

### Most populations were rescued, but overall performance varied during adaptation to corn

All evolving lines adapted to corn flour in a strikingly parallel manner, with dynamics during the first 20 transfers (one transfer is approximately one generation, so approximately 20 continuous generations) consistent with the classic U-shaped curve associated with evolutionary rescue (Fig. 3A-B). On average, populations showed a strong initial decline with about 15 dead adults/generation for the first 5-6 generations, then settled into a demographic bottleneck of a median of 12 live adults for several generations (hereafter ‘bottleneck’) before rebounding at a rate of about 25 adults/generation for a few generations. Surprisingly, only one population went extinct, but this enabled us to focus on testing the detailed dynamics and phenotypic drivers of population performance after successful rescue. For further analysis, for each evolving line, we estimated average population size (geometric mean of the number of adults/larvae) and coefficient of variation (CV; inverse of stability for adults/larvae) over ten generations, as detailed in the methods. We refer to a bin of ten generations as a ‘transfer bin’.

**Figure 3.**
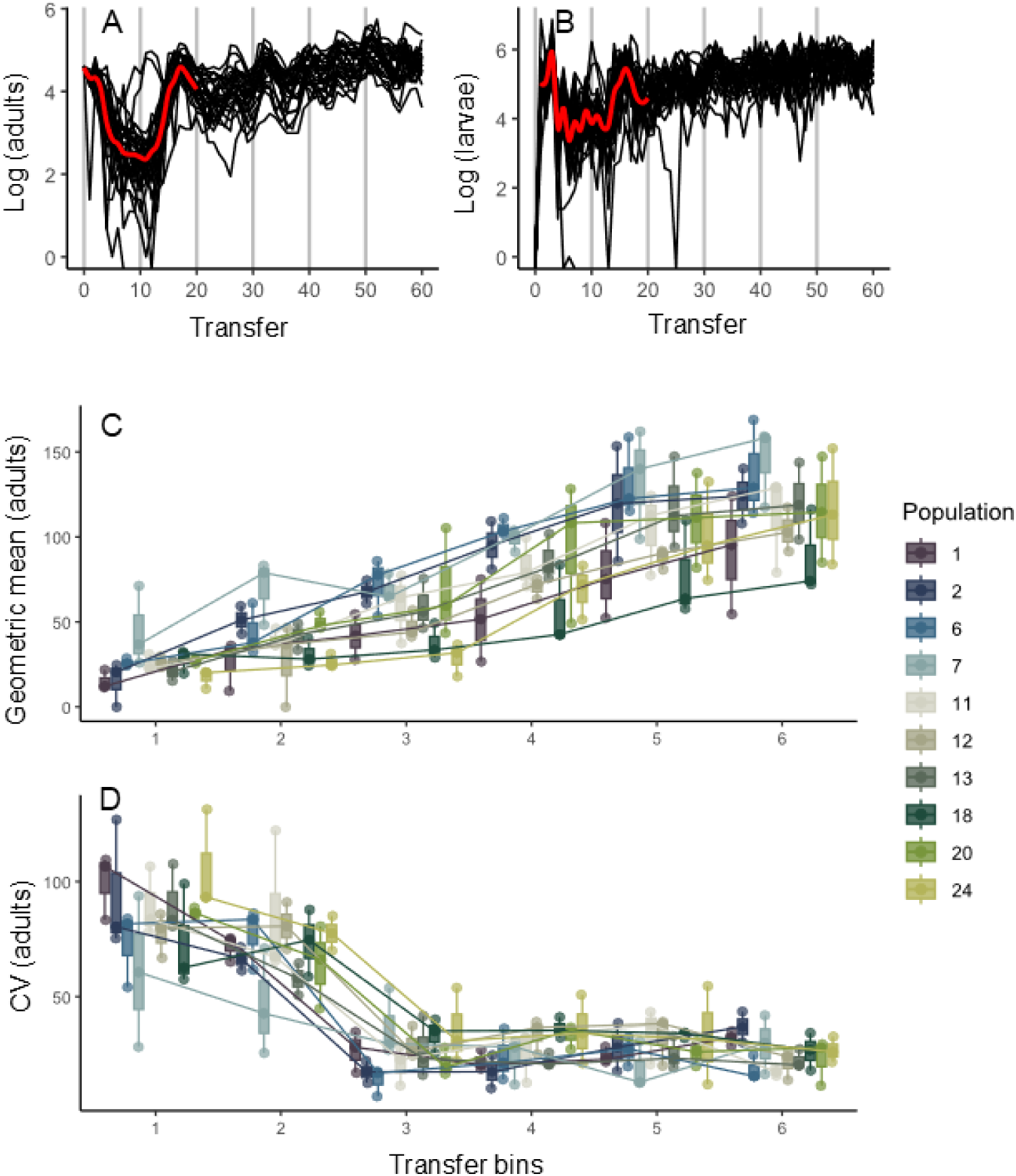
Overall demographic trends observed in corn-adapting lines. Time series of lines adapting to corn flour for (A) adults and (B) larvae, plotted on a natural log scale. The smoothed red curve (using loess) helps visualize the hallmark U-shaped curve of evolutionary rescue. Vertical grey lines represent transfer bins (10 generations per bin). (C) Average population size and (D) coefficient of variation (inverse of stability) across the adult time series for each transfer bin, for triplicates of each source population (three data points per boxplot; source populations are indicated by different colours). See Tables S4-S5 for ANOVA statistics of population performance across source populations.

We observed several signatures of adaptation – a gradual increase in both population size (Fig. S2A-B, Table S2) and stability over time (Fig. S2C-D, Table S3), with the average number of larvae staying consistent after approximately 40 generations, possibly indicating that the populations were close to the habitat carrying capacity soon after rescue. Despite the broad parallelism in the overall dynamics across independently evolving lineages, average population size varied significantly across source population identity for much of the experiment, for both life stages (Fig. 3C; Table S4). Adult population size broadly converged across source populations after generation 40, but this was not so for larvae (Table S4). In contrast, population stability did not vary by source population identity at any time (Fig. 3D, Table S5). Further, the performance rank order of individual selection lines was weakly consistent (over time) during rescue (until transfer bin 2, or generation 20; Table S6), after which bin-to-bin performance correlations strengthened, supporting the overall impact of source population identity on population performance.

### Population performance is most strongly associated with founder development rate

If specific founder traits drove differential population performance during and after rescue (Fig. 3C), we would see (a) significantly differentiated phenotypes across source populations (already reported above), and (b) a correlation between population performance and specific phenotypic trait values for founders. An initial principal component analysis showed that the first two principal components accounted for only 53% of the variance in the ancestral trait values. Therefore, we used original trait values to test for correlations with performance and demography (52), using the false discovery rate (FDR) method (53) to correct for multiple comparisons. After correction, we found two traits that were strongly positively correlated with both adult and larval population size (Fig. 4A-B) for several tens of generations during adaptation — offspring development rate and female longevity (Fig. 4C, also see correlation plots in Fig. S3). Aligning with our initial prediction, the strongest and longest-term association was between population size and the mean and variance of development rate for founding individuals. These were positively correlated with average adult population size throughout the experiment (Fig. 4C, Spearman’s rho ranging from 0.42–0.63, all coloured squares have p < 0.05 after correcting for multiple comparisons). Interestingly, the strength of both these correlations decreased during evolution (correlation coefficients reducing from 0.63 to 0.42 (mean), and from 0.69 to 0.43 (variance)). Mean female longevity in corn also stood out as a potential predictor of population performance, being strongly and positively correlated with the average number of larvae in the second half of our experiment (Spearman’s rho values ranging from 0.48–0.64; Fig. S3).

**Figure 4.**
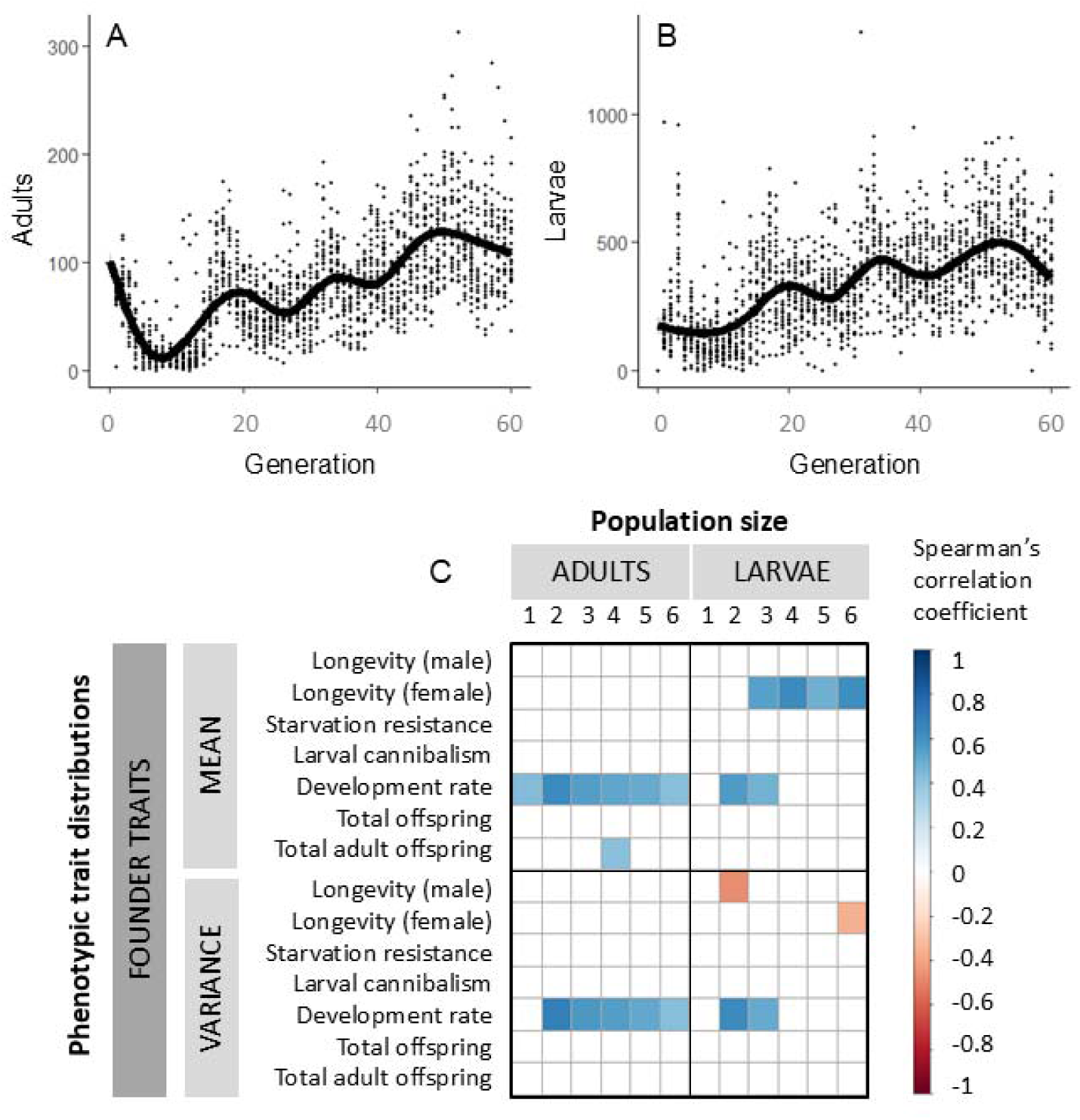
Ancestral phenotypic correlations for founder trait distributions with population size over time. Time series of (A) adults and (B) larvae combined across populations and replicates for visual reference. Black curves in both panels represent a smoothed fit of all data points per time point. (C) Matrices showing significant correlations (blue: positive, red: negative) corrected for multiple comparisons between ancestral phenotypic traits (means or variances) and the geometric mean of the number of adults and larvae evolving in corn per 10-generation bin (bins are numbered sequentially from 1-6), computed across lines. Only statistically significant Spearman’s rho correlation coefficients after correction for multiple comparisons are shown. Examples of raw correlation plots for development rate and female longevity are shown in Fig. S3.

### Founder traits do not predict specific features of population dynamics during or after rescue, and rescue demography does not predict later population dynamics

Since most of our tested lines underwent successful evolutionary rescue, we focused on across-line variation in the demographic decline, bottleneck and recovery during successful rescue. Given the surprising long-term association of ancestral trait variation with population performance for up to several tens of generations, we also assessed whether founder traits influenced demographic declines and rebounds following rescue. Doing so helped us disentangle two possibilities – (a) founder traits drove population dynamics only during rescue, after which population ranks stayed consistent, leading to apparent (indirect) long-term correlations between founder traits and population performance, and (b) ancestral phenotypic variation directly drove population performance throughout the experiment, via continued trait evolution, eco-evolutionary feedbacks or changing trait correlations. The second possibility could additionally explain the appearance of ancestral trait correlations with population size after 20 generations of evolution (e.g., as observed for female longevity).

In contrast to our results with the 10-transfer bins (Fig. 4), we observed almost no correlations between ancestral traits and specific features of population dynamics during and after rescue (Fig. 5; individual piecewise fits shown in Fig. S4). The only significant correlations after correcting for multiple comparisons involved total offspring and female longevity. For instance, the first and third recovery started later in populations with higher mean total offspring (Spearman’s rho=0.53 and 0.31 respectively, both P < 0.05). The start of the third recovery also correlated negatively with variance in female longevity in corn (Spearman’s rho=-0.534, P < 0.05). Importantly, the rates and timings of specific rescue demographic events did not correlate with post-rescue demographic events (all p > 0.05). For instance, contrary to our expectation, populations that declined more slowly or recovered faster from the bottleneck did not consistently perform better than other populations.

**Figure 5.**
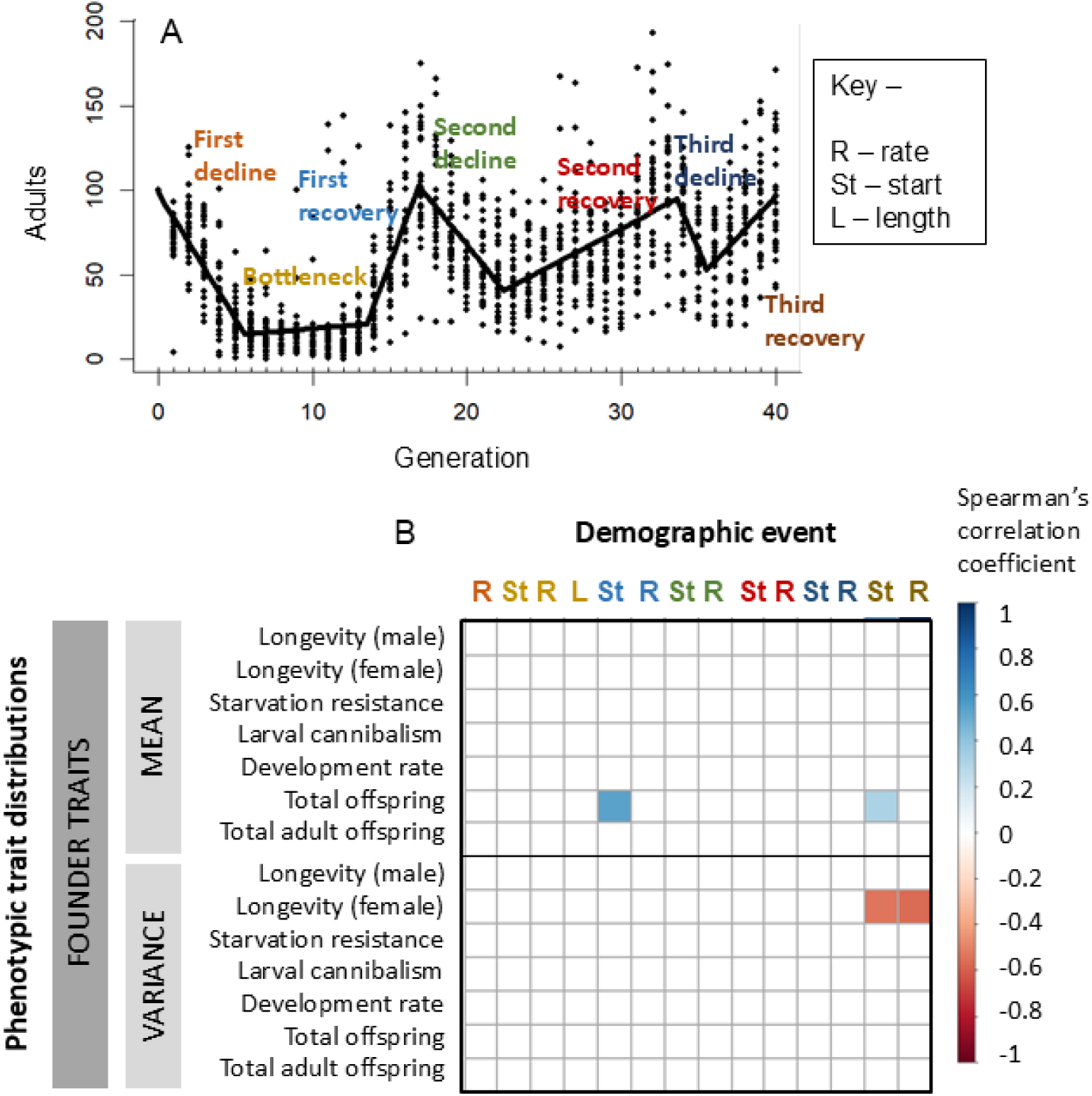
Ancestral phenotypic correlations for founder trait distributions with individual demographic parameters over time. (A) Time series of the adult population size, with the fitted global model combined across populations and replicates for visual reference, highlighting our identification of individual demographic events during and after the initial rescue. (B) Matrices showing significant correlations corrected for multiple comparisons (blue – positive, red - negative) between the individual demographic events with ancestral phenotypic traits (means or variances), computed across lines. Only statistically significant Spearman’s rho correlation coefficients after correction for multiple comparisons are shown. ‘Rates’ are extracted slopes from models fit individually to each evolved line; ‘Starts’ indicate the fitted breakpoint where one event ends and the other begins; and ‘Length’ refers to the bottleneck length preceding the rescue event.

Overall, these results suggest that while specific ancestral trait values had broad and consistent effects on population dynamics during corn adaptation across lines, they had weak predictive power with respect to specific demographic changes during and after rescue. Further, specific demographic phases during the evolutionary rescue itself were poor predictors of long-term population dynamics.

### Single-generation trait distribution shifts are typically adaptive but have poor predictive power for rescue and later dynamics

Given strong selection in the new corn habitat, we expected to find immediate and large changes in trait distributions within a single generation, which could influence population dynamics. Hence, we measured all the founder traits reported above (except longevity, due to the exceptionally long duration of this assay) for individuals approximating the first generation of each population. To do this, we allowed wheat-developed stock females to oviposit in corn, and then measured phenotypes for the resulting corn-developed adults (mimicking the 1^st^-generation offspring of the founders). Note that we did not use individuals from the evolving selection lines, as doing so would disrupt the experimental evolution, and it is unlikely that replicate lines from a single population would diverge significantly within a single generation. Thus, these measurements from source populations exposed to one generation in corn fairly approximate phenotypic shifts that likely occurred within a single generation in our selection lines, driven by single-generation mortality and/or plasticity.

Strikingly, all measured trait distributions changed significantly after a single generation in corn, with four of five traits showing adaptive shifts (Fig. S5; Table S7; note significant resource x source population interactions). Corn-reared (generation 1) females produced on average 1 more adult offspring than wheat-reared founders (all populations combined, GLM, total adult offspring ∼ development resource, p<0.001). These offspring also developed faster (p_development resource_ <0.001), even after accounting for offspring density in the assay plate (binomial GLM, proportion of pupae or adults at week 4 ∼ development resource x density, p_development resource_ < 0.01, other p values > 0.05). In contrast, average lifespan under starvation decreased in generation 1 compared to founders. Finally, larvae produced in generation 1 did not show higher cannibalism, which is adaptive in stressful environments (54). Notably, ancestral fitness in wheat was a poor predictor of immediate fitness in corn: the relative fitness values of founder vs. generation 1 beetles from a given population were not correlated for any of the measured traits (Spearman correlation ranks in Table S8, Fig. S6). In summary, development rate and the number of offspring (both total and adult) changed adaptively within a single generation in corn.

Given the broadly adaptive trait distributions in generation 1 individuals, we expected to find strong correlations with rescue dynamics and longer-term population performance. However, the variance in reproductive fitness at generation 1 was positively correlated with both adult and larval population size for 2-3 transfer bins only towards the end of our experiment, after about 30 generations of no correlation (Fig. S7; Spearman’s rho estimates ranging from 0.38–0.61). Other traits showed sporadic (also late) correlations (Fig. S7). Like founder traits, generation 1 traits were also typically uncorrelated with detailed population dynamics during and after evolutionary rescue (all p > 0.05, except the mean of starvation resistance and the start of the third recovery, Spearman’s rho estimate = 0.41). Thus, despite adaptive shifts in key traits such as offspring development rate and total offspring (Fig. S5), trait distributions after one generation in corn had poor predictive power for population performance during rescue, but showed some correlations with overall population performance in the later phase of population growth.

### Development rate and reproductive fitness increased across 70 generations

Our correlative analyses above indicated a major role for development rate in mediating performance in corn (Fig. 4). To directly test whether development rate also evolved as a result of selection, we measured development rate for each evolved line after 70 generations of evolution. Since total offspring and total adult offspring were also correlated with average population size (albeit at a later stage), we report them below as well. All three traits had increased significantly at generation 70, relative to both the founding individuals and generation 1 (Fig. 6A; Table S9). Beetle offspring in the evolved lines were overall 11 times and 4 times more likely to have successfully pupated in 4 weeks (i.e. had higher offspring development rate) relative to the (founding) wheat-developed ancestors and (generation 1) corn-developed individuals (top panel in Fig. 6A; Table S9). Further, females from evolved lines produced 4 more offspring (at all life stages) in corn relative to their wheat-developed ancestors, with a corresponding increase of 2 adult offspring (middle and bottom panels in Fig. 6A respectively; Table S9). However, the evolved development rate was not correlated with the ancestral development rate (Spearman’s rank correlation coefficient with founder and generation 1 development rate = 0.24 and 0.20; p > 0.05 for both), nor with population performance in the last 20 generations (Spearman’s rank correlations corrected for multiple comparisons, p>0.05). Interestingly, populations that recovered faster during rescue (rate of first recovery in Fig. 5A) had lower variance in evolved development rate after more than 50 generations (Fig. S8), indicating that rapid rescue may erode trait variation in the long term.

**Figure 6.**
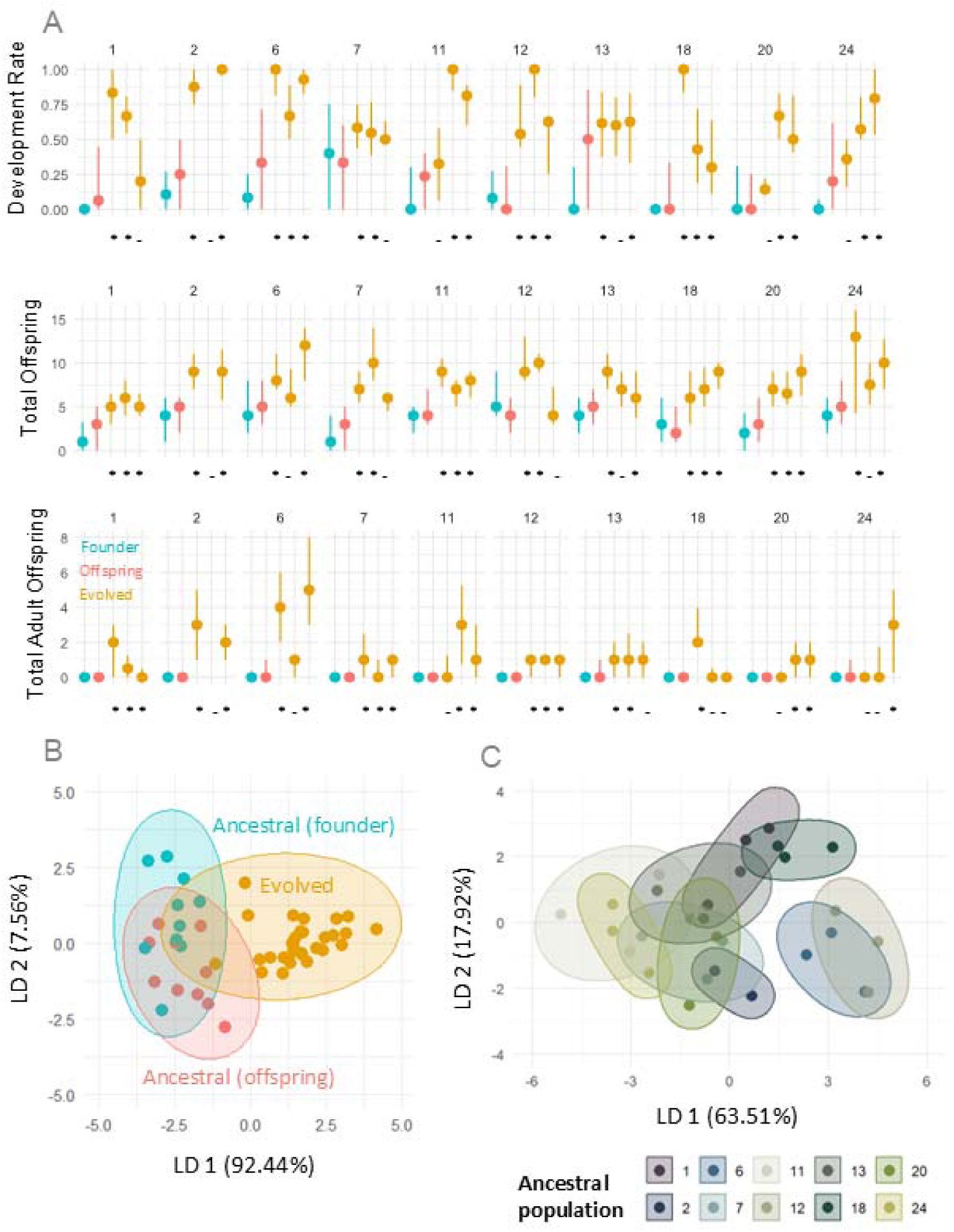
Ancestral and evolved development rate and reproductive fitness after 70 generation cycles of evolution. (A) The y-axes of the top, middle and bottom row of panels show development rate, total offspring and total adult offspring, respectively, for different populations. The x-axis shows trait values for ancestors developed in wheat (mimicking founding individuals; “founder”, blue), ancestors developed in corn (mimicking first-generation offspring; “offspring”, red); and three evolved replicates (yellow). Points indicate the median of the trait distribution and lines represent the interquartile range. Data shown for founders and offspring are identical to that in Fig. 2 and Fig. S4, replotted here for easy comparison. All significant differences are reported in Table S9 (between the evolved traits vs. the founder and offspring traits). Significant differences are represented as asterisks below each evolved trait distribution relative to the red offspring distribution within a panel; dashes indicate non-significant comparisons. Comparisons between the blue founder and red offspring distributions are shown in Fig. S5. Sample sizes per distribution range from 25 to 40 females. (B, C) Plots show the first two linear discriminants that group population phenotypes based on (B) ancestral founder and offspring traits or for populations evolved in corn for 70 generations (triplicate lines per source population), or (C) source population identity (see legend) for evolved traits. Data ellipses around the points are to visually highlight groupings only.

Finally, linear discriminant analysis revealed clear separation between the phenotypic space occupied by founders, our generation 1 approximation from the source populations, and the evolving selection lines at generation 70 based on the measured traits shown in Fig. 6A (Fig. 6B; PERMANOVA using Bray-Curtis distance, F = 51.243, R^2^ = 0.69, p = 0.001). A linear discriminant analysis of only the evolved trait distributions (at generation 70) did not indicate clustering by source population (Fig. 6C; PERMANOVA using Bray-Curtis distance, F = 1.6123, R^2^ = 0.056, p = 0.192). Thus, the strong effect of source population identity on long-term population dynamics did not shape evolved values of adaptive traits.

## DISCUSSION

We used comprehensive census data for 60 generations of experimental evolution in a harsh novel environment to determine which founding traits drove population dynamics during and after evolutionary rescue. Evolutionary rescue occurred within 20 generations, leading to large parallel increases in population size and stability. Development rate of founder offspring stood out as a strong and consistent driver of adaptation, with mean and variance in this trait being positively correlated with population performance (average population size). This trait also showed the highest initial variation between source populations compared to other fitness-related traits, potentially indicating past divergent selection. Development rate also increased dramatically within a single generation of exposure to corn (potentially reflecting both selective and plasticity effects), as well as over 70 generations of evolution. Finally, the negative association between the speed of evolutionary rescue and the variance in evolved development rate indicates surprisingly long-term potential effects of rescue dynamics on trait evolution. Our results complement prior work showing faster development rate associated with evolutionary rescue and adaptation to corn, along with other traits such as larval preference for corn and fecundity (31). A meta-analysis of arthropod studies further shows that juvenile development rate is especially important for thermal adaptation (55), suggesting that this trait may facilitate insect adaptation in several contexts. We also found a strong effect of female longevity on population size, likely reflecting increased lifetime reproductive output per female under the continuous generation cycles maintained in our experiment. Thus, our study provides an empirical example of specific traits consistently driving adaptation to a suboptimal habitat in populations founded by diverse source populations, across both single-generation and longer-term change. By using an extensive long-term dataset under controlled laboratory conditions and focusing on the detailed population dynamics during and after rescue, our work builds on prior studies demonstrating how adaptation is affected by intraspecific variation (56), single-generation trait changes (57), and both factors acting together (58).

Our results also indicate that the role of ancestral trait variation is dynamic. For instance, the impact of ancestral development rate was strongest during rescue and declined afterwards, during which correlations between population performance and other traits emerged. Therefore, we suggest that development rate was under selection more strongly in the early generations of the experiments, whereas other traits — such as reproductive fitness (offspring per female) — faced such selection later. Indeed, despite evidence suggesting strong selection on development rate and its role in driving adaptation, the evolved development rate at generation 70 was not correlated with late population performance. Nonetheless, it is possible that population performance at any given time may be strongly correlated with contemporaneous development rate. Overall, we suggest that distinct selection pressures likely operated across the course of the experiment, driving different correlations between various traits and population performance over time.

In contrast to the consistent positive correlations with overall population performance, ancestral trait distributions were not at all associated with specific demographic events during or after evolutionary rescue. In fact, none of the factors we tested — mean or variance in founder traits, mean or variance in traits of generation 1, and demographic events during early stages of rescue — were correlated with any aspect of rescue dynamics. This was surprising because we had predicted rescue rate and bottleneck duration to be the primary demographic events driving overall population performance, especially given the large amount of research generally linking demographic processes to evolutionary rescue (12, 13, 28). Could this lack of association in our experiment arise because the rescue dynamics were too similar across populations? This is not the case: the length of the initial population size bottleneck ranged from 4 to 11 generations across lines, and the slope of the subsequent recovery varied 18-fold across the fastest vs. slowest populations. Perhaps the populations were prone to high levels of demographic stochasticity during rescue, obscuring any effects of founder traits. In the end, it remains unclear what factors determined the speed and dynamics of evolutionary rescue in our study. Prior work also found that founding genetic diversity was not a good predictor of the rate of initial population decline, although it was strongly associated with the probability and magnitude of rescue in flour beetles (31). Interestingly, in our current work, early demographic events also did not correlate with later dynamics, suggesting that the rescue event itself (e.g., the bottleneck length or population size decline and recovery rates) did not contribute to shaping later dynamics. We speculate that this is because once rescue occurs, population dynamics are primarily driven by the strong negative density dependence typical of *T. castaneum*, which is relatively invariant across our source populations (59). Further work on other species with different mechanisms of population regulation may reveal whether the lack of longer-term effects of rescue dynamics is a general outcome, or specific to the particular life history of each species.

Could the highly parallel phenotypic shifts and population dynamics that we observed tell us something about the underlying genetic changes? Parallel trait evolution is found frequently, with many potential mechanisms and underlying processes (60, 61). We discuss two possibilities, differentiated by their timescales. The parallelism may be driven by strong initial maladaptation, evident from the large crash in population size. Such strong declines are broadly predicted to facilitate genetic and phenotypic parallelism (19), especially for sexually reproducing species with high standing genetic variation (62, 63). Parallel adaptation in such cases is likely driven by a few large effect beneficial alleles (19, 30), since the declining strength of selection in shrinking populations is unlikely to allow fixation of small-effect alleles during rescue (64). An alternative possibility is that parallel phenotypic evolution occurred after rescue. Given the timescale of our experiment, traits in all populations could slowly approach the optimum or maximal possible values in corn, even with multiple different small-effect alleles being fixed. If we had measured trait values soon after rescue, we may have been able to differentiate between these two possibilities. Note that despite the broad phenotypic convergence, our populations may have adapted via distinct genetic pathways (65) with different rates of adaptation, as observed in laboratory experimental populations founded with closely related yeast genotypes (66). This is more probable if the focal traits under selection are polygenic — likely true in this case — because initially different source populations could then more easily find diverse genetic routes to the fitness optimum (67–69). Yet another possibility is that some of the parallel dynamics — in particular, the second population decline — may have resulted from environmental forcing, e.g., changes in the texture of the corn flour used to impose selection (see Methods). However, since the correlation of performance with development rate remains consistent before and after this event, this seems unlikely to strongly alter the primary underlying evolutionary processes. Finally, we speculate that the strong and parallel correlations between population performance and ancestral traits may reflect a major role for standing genetic variation rather than *de novo* mutations in driving adaptation in our study (62, 63). In general, standing genetic variation is known to facilitate evolutionary rescue (8, 16, 20, 23, 31, 33). Identifying the genetic basis of functional trait variation as well as adaptation in these populations will thus allow us to understand the mechanisms underlying the remarkably parallel trait evolution during rescue and beyond.

An important result from our work is the dramatic shift in trait distributions between founders and their offspring within a single generation. Although we cannot discern whether the trait changes reflect selective mortality or plasticity (or a combination of both), the shifts were clearly adaptive, with the sole exception of starvation resistance. This pattern echoes findings from reciprocal transplant studies with plants: environmentally induced phenotypes (as in our ‘offspring’ generation 1) typically align with the direction of locally adapted populations (70). Adaptive plasticity can facilitate adaptive evolution on short time scales (43), and previous work has discussed the helpful role of plasticity in ‘buying time’ for adaptive evolution to occur prior to evolutionary rescue (41, 44). Importantly, trait distributions corresponding to the founder beetles (mimicking, e.g., individuals that completed their larval development elsewhere, then dispersed to a new resource patch) vs. their offspring (the first generation to complete their entire development in the new resource) were differentially correlated with population performance. *A priori*, we expected that performance in the new habitat would correlate more strongly with offspring trait distributions because these should be more relevant to the rescue event (in the new habitat) than the founder trait distributions. On the contrary, we observe a strong and long-lasting impact of founder traits (offspring development rate and female longevity), and only a late impact of offspring traits (reproductive fitness). Although we are not aware of other comparable experimental studies, prior analyses broadly examining the impact of parental effects on population dynamics and adaptation have also typically found moderate to strong parental impacts under diverse experimental conditions (47–50). Our results thus complement these prior studies and demonstrate the importance of parental effects on adaptation. We see very strong correlations between population performance and offspring trait values appearing as late as after 40 generations of evolution. We suggest that this may be because these phenotypic traits have weaker effects on initial performance in corn, impacting population performance only after rescue (also known as delayed life-history effects, or carry over effects) (71). Delayed life history effects are not rare (47, 72–74), but the magnitude of the time lag that we observed (40 generations) is much higher than documented earlier. Note that the lines were propagated in overlapping generations, so many founding individuals likely lived beyond the 1^st^ generation cycle and presumably continued producing offspring across multiple generations. However, the lag cannot be explained by exceptionally long-lived founders: 40 generations corresponds to about 1100 days, far exceeding the lifespan of any founder individual from any population in our dataset (about 270 days).

The surprisingly low frequency of extinction in our populations (1 in 30) begs the question of whether our populations underwent evolutionary rescue or simply adaptation. We believe the dynamics indeed represent evolutionary rescue, for the following reasons. One of the hallmarks of evolutionary rescue is a U-shaped curve in population size over time, which is very clearly observed in our census data. Prior work has reported U-shaped trajectories consistent specifically with evolutionary rescue of *T. castaneum* populations experimentally evolved in corn flour (13, 22, 31). All these studies reported multiple extinctions in lines adapting to corn flour, suggesting that, in general, *T. castaneum* is strongly challenged by corn (75). Populations in our study reached a median bottleneck population size of 12 adults (± 5.17 SD), which is consistent with the bottleneck sizes reached in prior work where extinction was common (31). It is likely that since our populations were wild-collected, they had greater genetic diversity (compared to the laboratory-inbred lines used in prior work), potentially reducing extinction probability. It is also likely that the maintenance of continuous generations in our experiment (which allowed adults to reproduce for a long time) buffered populations against extinction. This would also explain how offspring phenotypic variation was preserved in the population for as long as 40 generations, allowing for a late, but strong, impact on population performance. If true, this suggests that species that reproduce continuously may show counter-intuitive adaptive trajectories during and after rescue, as observed in our study. We suggest more empirical work comparing discrete vs continuously reproducing populations or species for a more nuanced understanding of extinction and rescue.

In summary, we used comprehensive datasets for a wide array of ancestral fitness-relevant phenotypic traits and a 70 generation long experimental evolution system to establish a central role for source population identity during and after evolutionary rescue. Further, we identified founder development rate and female longevity as key contributors to rescue, with most populations showing large adaptive shifts in development rate within a generation as well as across 70 generations in corn. While several studies have examined different aspects of our work in isolation, we were able to study all of them together in a single experimental system. Many more empirical studies with diverse species and methods are needed to catch up with the burgeoning theory in the rescue literature, and also to assess the generality of our findings. Nonetheless, our study already has several implications for our ability to predict evolutionary change in several contexts (61). Our results suggest that some pest populations may be especially difficult to eradicate because fast juvenile development may allow rapid evolutionary rescue under stressors such as novel suboptimal resources or pesticides, facilitating convergence on fitness peaks. Conversely, local populations with a high intrinsic capacity for rapid adaptation under stress (e.g., with high development rates and reproductive fitness) may be best suited for species conservation via re-introduction efforts. Our results also indicate that detailed information on rescue dynamics may not be necessary to predict future evolution, although much more work is needed to rigorously test this idea. Together with a growing body of related work, we hope that our study will aid the prediction of evolutionary responses to climate change and other anthropogenic stressors (76).

## METHODS

### Collection and maintenance of source populations and selection lines

We used 10 wild-collected populations of *T. castaneum* from different locations in India (50-100 individuals per source population, Fig. 1A) and established large laboratory stocks of these populations in plastic boxes (diameter 21 cm, height 8.5 cm) with holes in the lid for ventilation. To propagate stocks, we allowed approximately 2000 adults (1-2 weeks post-eclosion) to oviposit in 750g whole wheat flour for 7 days, following a discrete generation cycle, after which we allowed offspring to develop for 28 days. This regime ensured at least 2000 individuals every generation, facilitating maintenance of genetic diversity. Stocks were acclimatized to laboratory conditions in dark Percival incubators at 32°C (±1°C) for 2-3 years (20-30 generations) prior to initiating the selection lines. Note that populations are not numbered 1-10 consecutively; since they were drawn from a larger collection of source populations, they correspond with laboratory IDs.

To study evolutionary rescue, we used a highly suboptimal novel resource, corn flour. *T. castaneum* populations established in corn show low population growth rates (15), and are often driven to extinction (22). We established a total of 30 experimental lines in corn (10 source populations x 3 replicates each; Fig. 1B) founded with 100 pupae each from the respective source population stocks, which have a typical sex ratio of 1:1. In contrast with the stock maintenance, we propagated the selection lines in overlapping generations. We initiated lines in a staggered manner over 6 months, so that the three replicates of each source population were started on different days. We maintained all lines in 50 g (±0.5 g) corn flour in plastic boxes (6.7 cm diameter) with small holes in the lid for ventilation. We purchased flour from the same supplier throughout and documented any changes in the texture of the corn (Fig. S9). To kill any prior pest infestations, new flour packets were sterilized by freezing at –80°C for 24 hours. We maintained the lines in incubators as above.

Every 28 days (± 1 day) — corresponding to the average development time from egg to pupae in corn (i.e., the approximate generation time) — we replenished the resource by transferring populations to fresh flour. At each transfer, we censused all life stages except eggs (i.e., we counted dead and living larvae, pupae and adults), by sifting the contents of the box through meshes of different pore sizes (600-µm and 710-µm; Daigger), and discarding dead individuals. Eggs typically take 48 hours to hatch, so all eggs laid 2-3 days prior to the census were discarded along with the used flour. We heat-sterilized all equipment for a minimum of 10 minutes at 65°C to prevent cross-contamination.

### Assessing population performance

One line (replicate 2 of source population 2) went extinct within 5 generations and was excluded from all further analyses. We also excluded census data between generations 60-70 from subsequent analysis since we encountered many COVID19-related disruptions to the workflow during this time. We evaluated the overall population performance for each selection line from the census data for adults and larvae (the most abundant life-stages in the lines) by calculating the geometric and harmonic mean population size. The former is a natural scale for population size since populations grow multiplicatively, and the latter is an approximate estimate of effective population size (77) and accounts for population size fluctuations (78). Additionally, we estimated population stability by quantifying population size oscillations using the coefficient of variation (CV). We split the dynamics (and calculations) into bins of exactly ten successive generations (i.e., generations 1–10, 11–20, until generation 60) and evaluated population performance (average size and CV) per bin to minimize autocorrelation effects between each individual generation. The harmonic and geometric means were strongly correlated for both adults and larvae (Pearson’s correlation > 0.99), so we only report geometric means throughout. As mentioned above, each geometric mean and CV value is calculated using ten successive data points per evolving line. In addition, as a direct assay of adaptation we measured two important traits in each evolved line, as described in the next section.

### Measuring fitness-related phenotypic traits

We assayed the ten source populations for five fitness-related traits: adult longevity (lifespan after eclosion with access to food), adult starvation resistance (lifespan after eclosion without food), larval cannibalism (number of eggs cannibalized per larva per day), offspring development rate (proportion of eggs that developed to the pupal or adult stage in 4 weeks, hereafter ‘development rate’), and a snapshot of adult reproductive fitness (viable offspring produced per female – either total, or those that developed to adulthood) (Fig. 1; Fig. S1; SI methods). Our measures of reproductive fitness are defined this way to contrast with other phenotypes directly related to survival (adult longevity and starvation resistance), and other traits that are indirectly linked to either survival, reproduction, or both (cannibalism and offspring development rate). Our snapshot reproductive fitness measures implicitly include female fecundity, egg hatchability and survival to late larval stages within the assay, but we do not measure each of these individually.

We were not able to measure phenotypes of the founding individuals directly, since this would disrupt the experimental evolution. To mimic founder trait distributions (‘G0’, hereafter ‘founder phenotypes’), we allowed individuals from each of the ten source populations to develop in wheat, and then transferred them to corn to measure the phenotypes. This mimics the founding individuals’ developmental environments, in that they founded the populations in corn after stock maintenance in wheat. We also used an experimental proxy for the phenotypic distributions of the first generation of individuals within the selection lines that developed in corn (‘G1’, i.e., the offspring of the founders, hereafter ‘offspring phenotypes’ or ‘generation 1 phenotypes’), by measuring all the above phenotypes for stock individuals whose parents reproduced in corn, leading to the test individuals developing in corn too). Note that we did not measure these offspring phenotypes from the actual selection lines, as doing so would interfere with experimental evolution. Doing so allowed us to test the relative importance of the phenotypic distributions of the founders versus their offspring (with different development resources), giving us more insight into potential disconnects between ancestral variation and population dynamics. Finally, to test a hypothesis that arose during data analysis (discussed in Results), we re-measured two traits (development rate and reproductive fitness) after 70 generations of evolution of each evolving population (hereafter ‘evolved phenotypes’, from the actual experimental evolution setup). We did this by deriving smaller sub-lines from the evolving lines, which we then measured (SI methods).

### Model fitting to adult population dynamics using piecewise linear models

To determine distinct stages of evolutionary rescue, we divided the population dynamic trajectories to reflect population recoveries, declines and bottlenecks. To clarify, we use ‘bottleneck’ to only refer to the period of consistently low population sizes after the initial decline and before the first recovery (see Fig. 5). We first fit a piecewise model to the combined data from all populations. We only used data until 40 generations (or generation cycles) for this analysis since the lines showed more divergent dynamics after this time. After trying several alternate models with different numbers of breakpoints, we chose the simplest model that captured the most variance in the data, settling on a model with 6 breakpoints that captured 51.8% of variance in the combined dataset. The breakpoint parameters from this ‘global’ model were then used to fit the same model to each individual line to prevent overfitting and to enable easy comparison. The global model captured fundamental population processes such as change in population size and bottlenecks. For the individual model fits, 6 models captured >90% variation; 17 captured 80-90% variation, and 5 captured 60-80% of the variation in the respective population dynamics. We excluded two lines from further analysis, one for which the model captured <60% of variation (population 7 replicate 3), and one that went extinct at generation 5 (population 2 replicate 2). We extracted slopes and breakpoints to quantify the variation in dynamics for each individual line, then used these data for further analyses to test for correlations with ancestral traits and post-rescue performance.

### Data analysis and visualization

All analyses and plots were generated in R version 3.4.4 in RStudio version 1.0.143 on a Windows 11 system. Figures were created in Microsoft Powerpoint, GIMP and Affinity Photo. Statistical tests are reported directly in the main text or Supplementary Information legends for clarity. Packages used for analysis and visualisation were: ggplot2, gridExtra, paletteer, dplyr, tidyr, forcats, hrbrthemes, viridis, corrplot, RColorBrewer, survival, sjstats, segmented, broom, ggpubr, MASS, and vegan.

## ACKNOWLEDGEMENTS

We thank Pratibha Sanjenbam, Swastika Issar, Shazia Parveen and other members of the Adaptation lab for comments on the manuscript. We thank two anonymous reviewers for their exceptionally constructive and detailed comments on the manuscript.

## SOFTWARE AND PACKAGES

Auguie, B. and Antonov, A. (2017) gridExtra: Miscellaneous Functions for “Grid” Graphics. R Package Version 2.3. https://CRAN.R-project.org/package=gridExtra

Garnier, Simon, Ross, Noam, Rudis, Robert, Camargo, Pedro A, Sciaini, Marco, Scherer, Cédric (2024). viridis(Lite) - Colorblind-Friendly Color Maps for R. doi:10.5281/zenodo.4679423, viridis package version 0.6.5, https://sjmgarnier.github.io/viridis/.

Hvitfeldt E (2021). paletteer: Comprehensive Collection of Color Palettes. R package version 1.3.0, https://github.com/EmilHvitfeldt/paletteer.

Kassambara A (2023). *ggpubr: ‘ggplot2’ Based Publication Ready Plots*. R package version 0.6.0, https://rpkgs.datanovia.com/ggpubr/.

Lüdecke D (2024). *sjstats: Statistical Functions for Regression Models (Version 0.19.0)*. doi:10.5281/zenodo.1284472, https://CRAN.R-project.org/package=sjstats.

Muggeo VM (2008). “segmented: an R Package to Fit Regression Models with Broken-Line Relationships.” *R News*, **8**(1), 20–25. https://cran.r-project.org/doc/Rnews/.

Neuwirth, E. (2014) RColorBrewer: ColorBrewer Palettes. R Package Version 1.1-3. https://CRAN.R-project.org/package=RColorBrewer

Oksanen J, Simpson G, Blanchet F, Kindt R, Legendre P, Minchin P, O’Hara R, Solymos P, Stevens M, Szoecs E, Wagner H, Barbour M, Bedward M, Bolker B, Borcard D, Borman T, Carvalho G, Chirico M, De Caceres M, Durand S, Evangelista H, FitzJohn R, Friendly M, Furneaux B, Hannigan G, Hill M, Lahti L, McGlinn D, Ouellette M, Ribeiro Cunha E, Smith T, Stier A, Ter Braak C, Weedon J (2025). *vegan: Community Ecology Package*. R package version 2.7-0, https://github.com/vegandevs/vegan, https://vegandevs.github.io/vegan/.

R Core Team (2021). R: A language and environment for statistical computing. R Foundation for Statistical Computing, Vienna, Austria. URL https://www.R-project.org/.

Robinson D, Hayes A, Couch S (2024). *broom: Convert Statistical Objects into Tidy Tibbles*. R package version 1.0.7, https://github.com/tidymodels/broom, https://broom.tidymodels.org/.

RStudio Team (2020). RStudio: Integrated Development for R. RStudio, PBC, Boston, MA URL http://www.rstudio.com/.

Rudis B (2025). *hrbrthemes: Additional Themes, Theme Components and Utilities for ‘ggplot2’*. R package version 0.90, https://github.com/hrbrmstr/hrbrthemes.

Therneau T (2024). *A Package for Survival Analysis in R*. R package version 3.8-3, https://CRAN.R-project.org/package=survival.

Terry M. Therneau, Patricia M. Grambsch (2000). *Modeling Survival Data: Extending the Cox Model*. Springer, New York. ISBN 0-387-98784-3.

Venables WN, Ripley BD (2002). *Modern Applied Statistics with S*, Fourth edition. Springer, New York. ISBN 0-387-95457-0, https://www.stats.ox.ac.uk/pub/MASS4/.

Wickham H (2016). ggplot2: Elegant Graphics for Data Analysis. Springer-Verlag New York. ISBN 978-3-319-24277-4, https://ggplot2.tidyverse.org.

Wickham H, François R, Henry L, Müller K, Vaughan D (2023). dplyr: A Grammar of Data Manipulation. R package version 1.1.4, https://github.com/tidyverse/dplyr, https://dplyr.tidyverse.org.

Wickham H (2023). forcats: Tools for Working with Categorical Variables (Factors). R package version 1.0.0, https://github.com/tidyverse/forcats, https://forcats.tidyverse.org/.

Wickham H, Vaughan D, Girlich M (2024). tidyr: Tidy Messy Data. R package version 1.3.1, https://github.com/tidyverse/tidyr, https://tidyr.tidyverse.org.

Wei T, Simko V (2024). *R package ‘corrplot’: Visualization of a Correlation Matrix*. (Version 0.95), https://github.com/taiyun/corrplot.

