## Supplementary Information for "Founders predict trait evolution and population performance after evolutionary rescue in the red flour beetle"

**SUPPLEMENTARY METHODS**

**Measurement of individual phenotypic traits**

After initiating the selection lines, ancestral population stocks were switched to a lower maintenance regime. They were kept on an eight-week discrete generation cycle with at least 500 adults every generation. The flour was replaced and dead individuals were discarded at least once every generation. All stock populations were kept at 33°C (±0.5°C) in a dark incubator.

*Longevity*

Adult longevity is an important indicator of fitness, since all else being equal, longer-lived adults are more likely to leave more offspring. To estimate beetle longevity in isolation, we sexed and isolated stock pupae (~30–40 pupae per sex per population) in 1.5 mL Eppendorf tubes containing 1 g of corn flour, with a small hole in the lid for ventilation. Tubes were monitored twice a week for 10 months, and the day of death was noted when relevant. The flour in each tube was changed every 4 weeks to mimic the selection line protocol. Longevity was only measured for wheat-developed individuals that were transferred post-pupation either into wheat or corn; although we only used the dataset for longevity in corn for the analyses with the selection lines.

*Starvation resistance*

Starvation resistance was measured as the lifespan of newly eclosed adults under starvation and indicates the amount of resources (stored during the larval stage) available for reproduction and survival in a harsh environment with limited or suboptimal food. We collected pupae from ancestral stock populations that developed either in wheat or corn (to mimic the conditions for founders and their offspring respectively). We followed the regime used for the ancestral stock populations, viz, ~2000 adults approximately 1-2 weeks post eclosion were allowed to oviposit for 7 days in either wheat or corn, after which they were removed, and offspring were allowed to develop for at least 21 days, after which populations were checked every alternate day for pupae. Each pupa was individually placed in a single well of a 48-well Corning micro-plate (i.e., 48 pupae per sex per population) with no flour. We observed pupae every day in the morning and evening (~10 hours apart) and noted the time of eclosion and subsequent death. We calculated starvation resistance as the time from eclosion to death for each population, with larger values indicating greater starvation resistance.

*Cannibalism*

Egg cannibalism is an important source of nutrition in suboptimal habitats (Ho and Dawson 1966; Via et al 1999) and population cycling in *Tribolium* has been linked to larval cannibalism of eggs (Benoit et al 1998). Hence, we measured per capita larval egg cannibalism for each population. For the sake of completeness, we assayed larvae developed in either wheat or corn (both assayed in corn), but we note that cannibalism by wheat-developed larvae would not occur in our selection lines since we initiated populations with pupae. We generated test larvae by allowing 200 stock adults to oviposit in 150 g of either wheat or corn for 24 hours. After this, we removed the adults and allowed larvae to develop for 14 days. We housed five randomly-picked 2-week-old larvae with 20 randomly-chosen eggs (oviposited by 400 adults from the same population one day earlier) in 1 g of corn flour in 30 mm Petridishes (n=25 plates per population). We used corn flour that was pre-sifted through a sieve with a very small pore size, to ensure that flour particles were smaller than eggs, allowing eggs to be counted. We removed larvae after 24 hours and counted the remaining eggs in each dish. For each plate, we calculated per capita larval cannibalism rate as (number of eggs added — number of eggs surviving after 24 hours) divided by the number of larvae in the plate at the end of the assay (to account for rare instances of larval death during the assay). We conducted the assay with groups of larvae (rather than single larvae) to ensure that the number of eggs cannibalized in each plate would be sufficiently large for us to accurately estimate cannibalism rate within a 24-hour period (before eggs start hatching).

*Development rate and reproductive fitness*

In addition to measuring the ten ancestral populations, we also measured these two traits (development rate and reproductive fitness) for all 29 evolved lines at transfer 70. We did not include population dynamics between generation 60 and 70 since there were multiple changes in the workflow during this time due to COVID. To phenotype the lines, we created derived lines from the regular selection lines at the time of their regular scheduled census. We allowed all available adults to lay eggs in 50g of fresh unsifted flour for 48-72 hours. After this, we removed the adults and returned them to their respective containers. These eggs and were allowed to pupate in corn for before they were isolated for the assay described below.

Faster-developing offspring have a selective advantage under strong competition, as do females that produce more viable offspring in a given time period. Thus, we measured development rate and reproductive fitness in corn as fitness-relevant traits (for individuals developed in either wheat or corn, representing founders and offspring respectively). We segregated male and female stock pupae into 1.5 mL Eppendorf microcentrifuge tubes (4 pupae per sex per tube) with 1 g of wheat or corn flour for 14 days, allowing pupae to eclose and mature sexually to an age that corresponds with peak female fecundity. We then allowed a randomly picked virgin male and a female to mate for 48 hours in a fresh tube containing the respective development resource. We sorted sexes under the microscope after giving a short cold shock to reduce movement and enable observation of the male femoral sex glands. We allowed the females to oviposit alone for 24 hours in a 60 mm Petridish containing 5 g corn flour (n = 25-40 females). After removing the female, we allowed offspring to develop for 4 weeks, and counted the number and life stage of all offspring. At the 3-week and 4-week mark, we counted and removed newly-developed adult offspring to prevent interference with the development of their siblings. We extracted the following fitness components from these data: offspring development rate (fraction of pupae and adults out of all surviving offspring at week 4), total offspring (total number of individuals at week 4), and total adult offspring (total adult offspring developed at or before week 4). We reiterate here that ‘development rate’ refers to offspring development rate. Due to the design of the assay, each female constitutes a single replicate, and so offspring development rate may be influenced by density-dependent competition, which we test for later.

**SUPPLEMENTARY FIGURES**

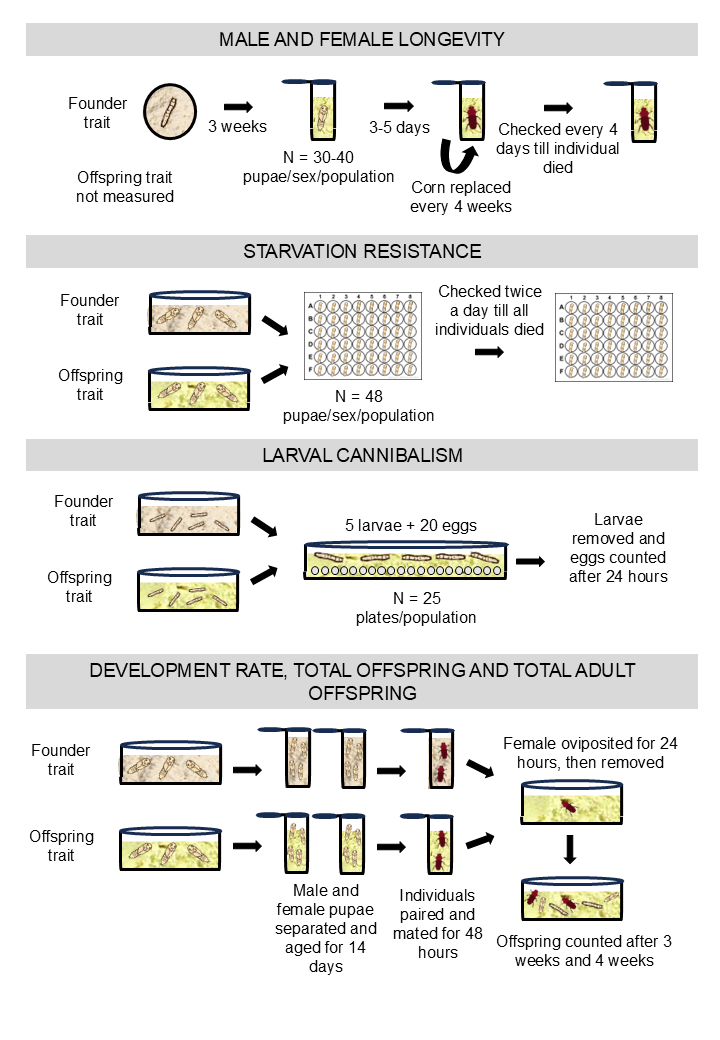
 **Figure S1.** Measurement of different phenotypes. Refer Supplementary Methods for details. Objects in the image are not to scale.

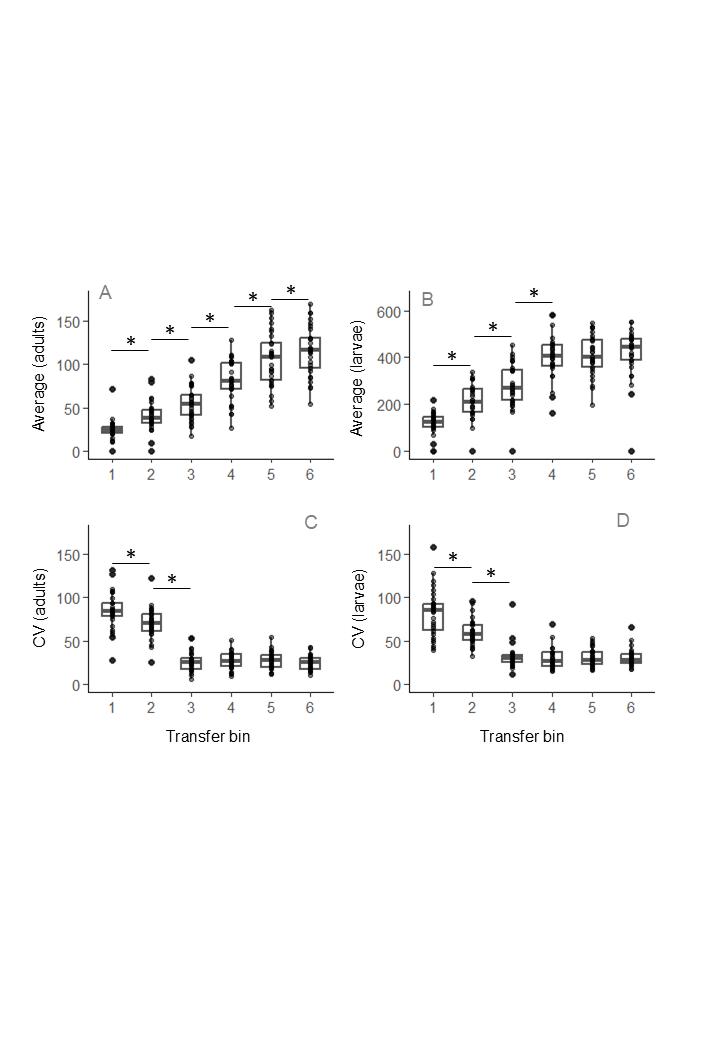
**Figure S2.** Population size per line (geometric mean over 10 generations or 1 transfer bin) for (C) adults and (D) larvae, and coefficient of variation (calculated per transfer bin) for (E) adults and (F) larvae. Significant changes in average population size and CV between consecutive transfer bins are indicated by an asterisk. Ten population transfers have been combined into 1 transfer bin to facilitate visualization of population trajectories.

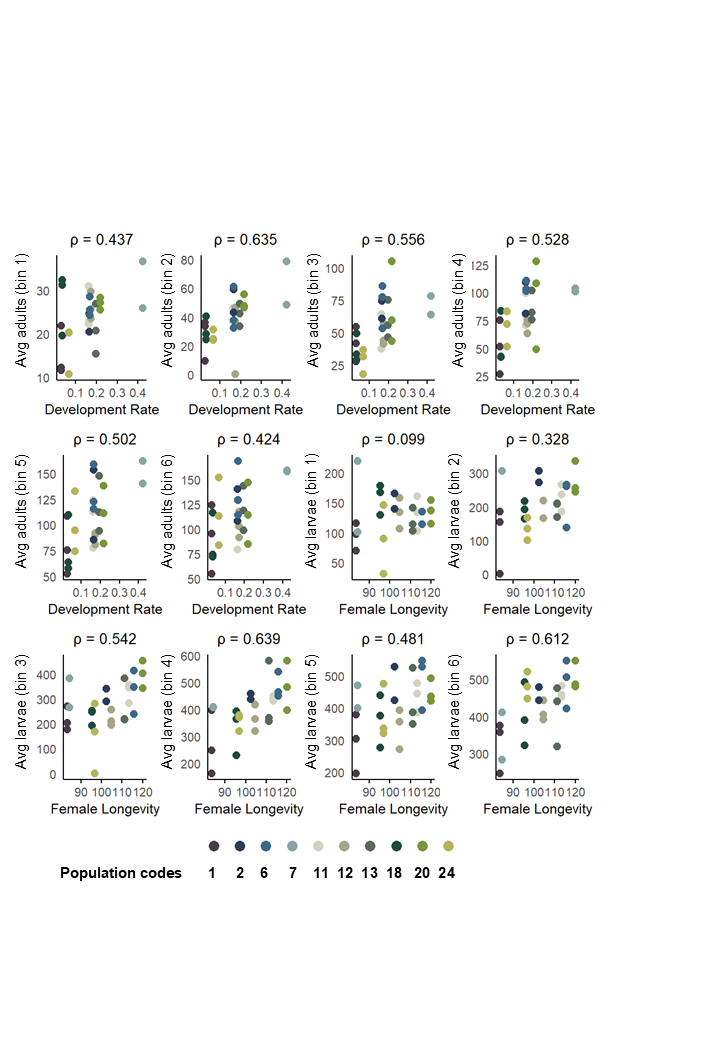
**Figure S3.** Correlation between key traits — ancestral development rate or female longevity (x-axis) — and average adult and larval population size (average population size in each transfer bin consisting of 10 generations each). Correlations such as these underlie the heatmap shown in Fig. 4C. Spearman’s rank correlation coefficients are given for each panel; statistical significance is indicated in Fig 4C, after correcting for multiple comparisons. For these correlations, we removed one replicate of population 2 that went extinct, and one replicate from population 7 that did not follow the average decline-recovery trends (see Fig. S4 below for individual trajectories).

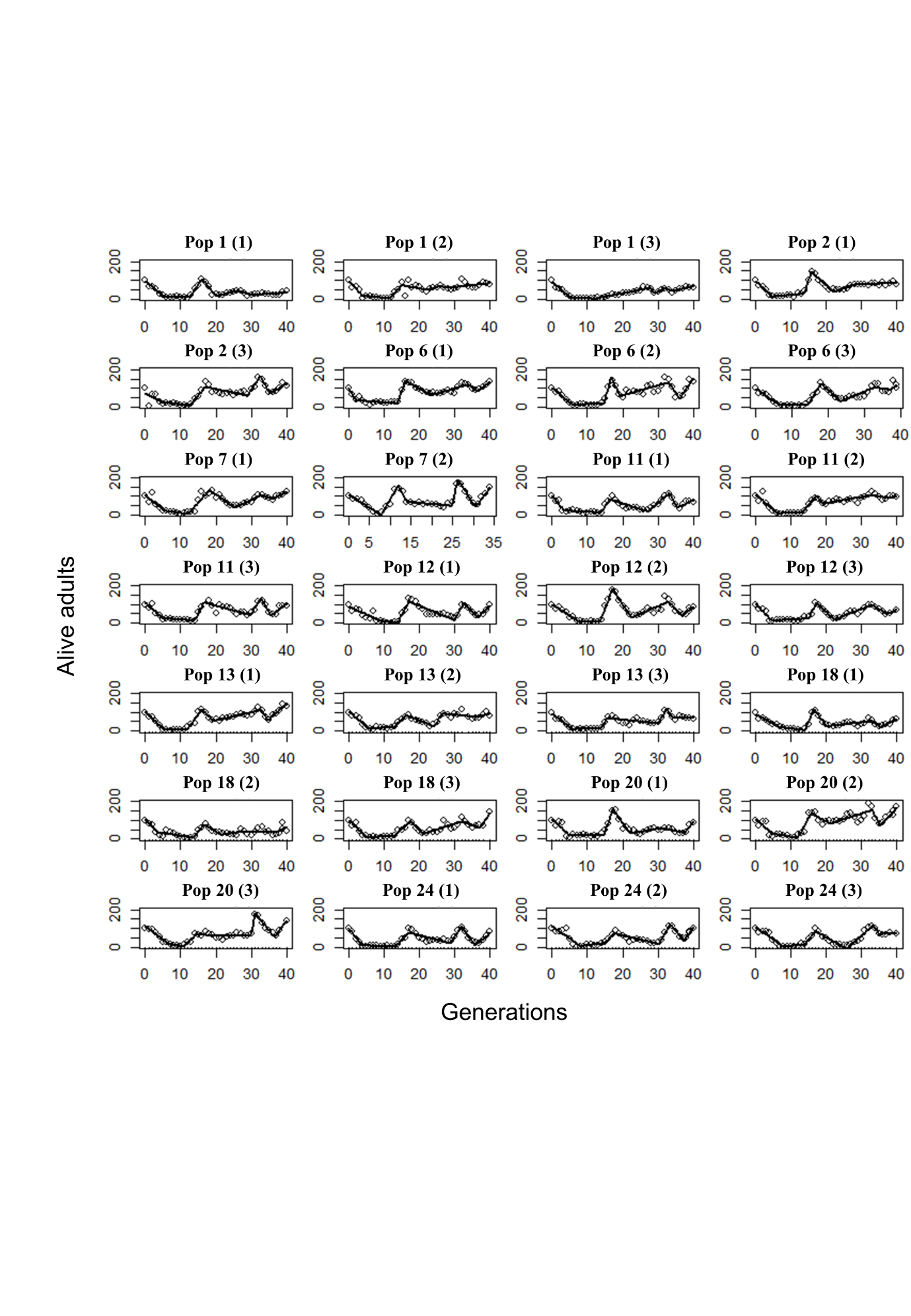

**Figure S4. Piecewise linear models fit to individual selection line dynamics**. Model fits range from explaining 60% to 95% of the variation in demographic time series. Points represent the data, and the lines connect adjacent points for ease of visualization. Replicate lines of a given source population are distinguished by the numbers in parentheses.

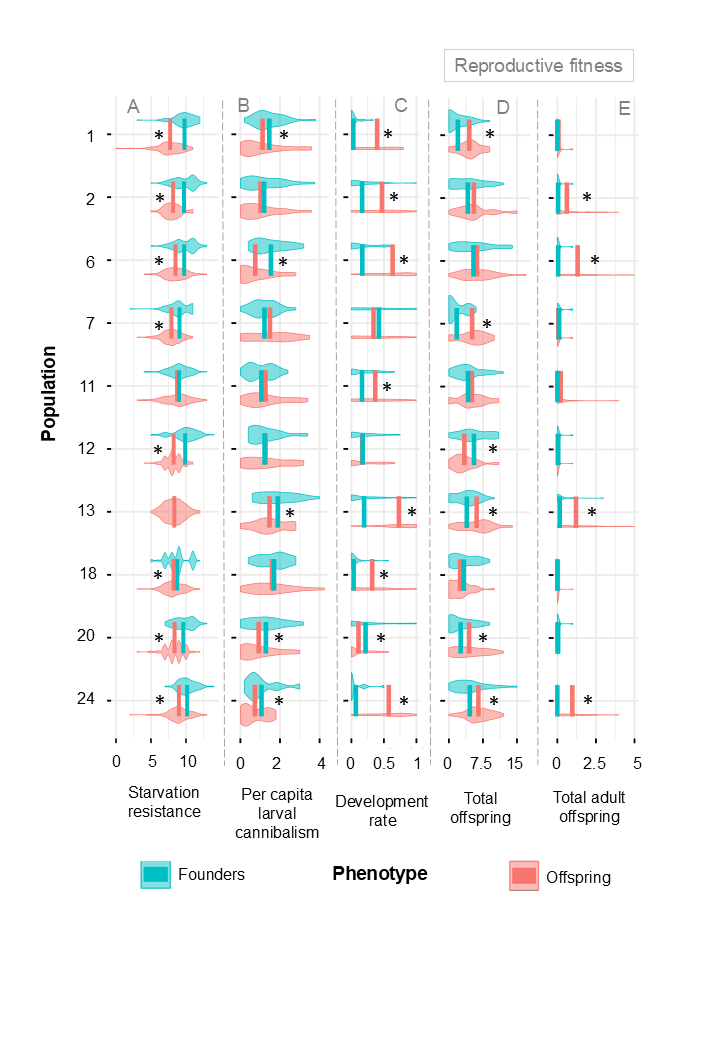

**Figure S5. Change in ancestral phenotypic traits after developing in wheat vs corn, representing the phenotypic trait distributions of the founders vs their offspring (generation 1) respectively.** Phenotypic trait distributions for each source population (y-axis) for different fitness-related traits (x-axis), with solid lines comparing distribution means for the same population across development resource. (A) Starvation resistance (n = 48 per population per resource); (B) larval cannibalism (n = 25 per population per resource), (C) offspring development rate (n = 25-40 females per population per resource) and (D-E) measures of reproductive fitness per female (n = 25-40 females per population per resource) such as (D) number of total offspring, and (E) number of adult offspring. The last two traits are what we jointly refer to as ‘reproductive fitness’. We were not able to measure the generation 1 distribution of adult longevity, so it is not included here. Founder starvation resistance assays for population 13 in panel A were lost to an accident, so this comparison is not included in the statistics. One extreme data point each is not shown in panels D (Population 12, wheat, 42 total offspring) and E (Population 6, corn, 10 total adult offspring) for ease of visualization, but the data were included in the reported statistics. Asterisks represent statistically significant differences (p<0.05) for the following tests in the respective panels: (A, D, E) LM and (B, C) GLM with binomially distributed errors.

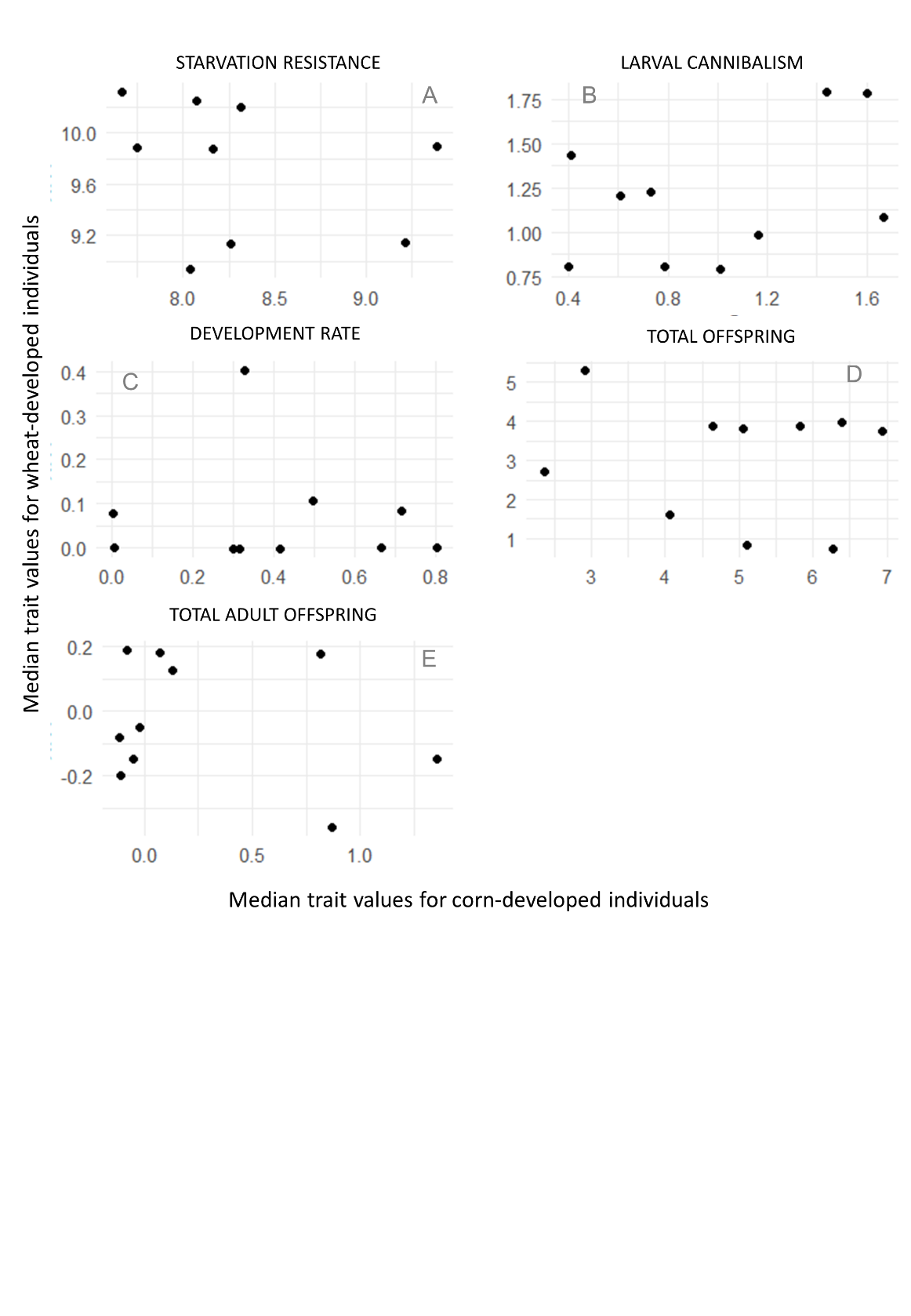

**Figure S6. Scatter plots for ancestral median trait values for individuals developed in wheat vs. corn.** Panels show data for (A) starvation resistance, (B) larval cannibalism, (C) development rate, (D) total offspring and (E) total adult offspring for individuals from the ancestral population developed in corn (x-axis) vs wheat (y-axis).

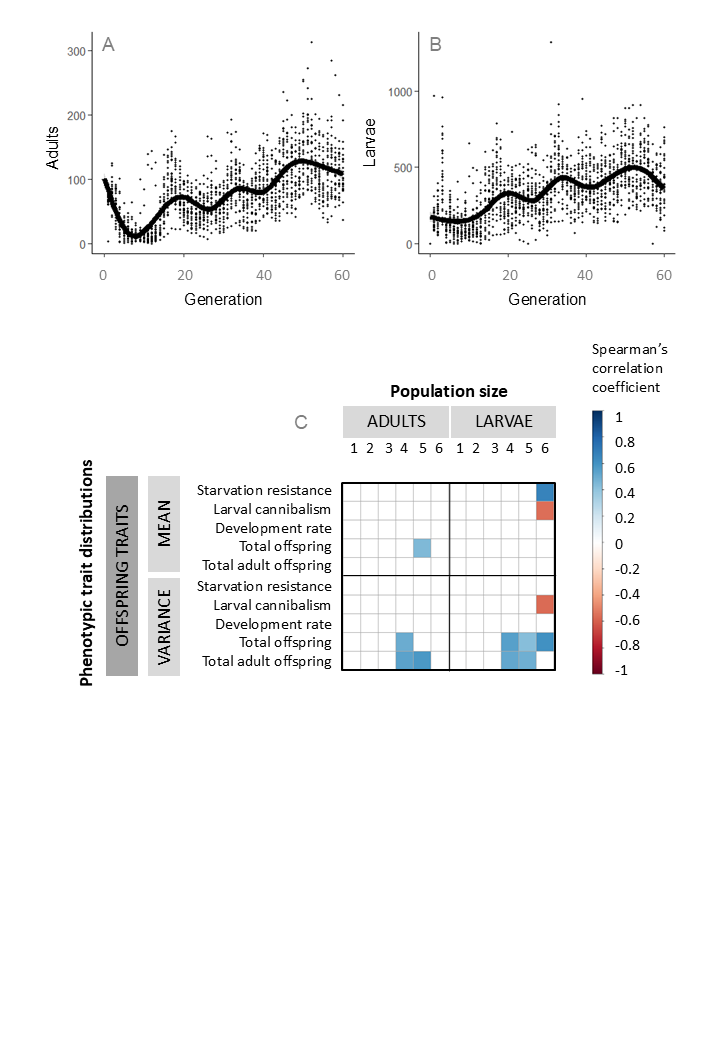
**Figure S7.** **Ancestral phenotypic correlations for offspring trait distributions (means and variances) with population size over time.** Time series of (A) adults and (B) larvae combined across populations and replicates for visual reference. Black curves in both panels represent a smoothened fit of all data points per time point. (C) Matrices showing significant correlations (blue: positive, red: negative) corrected for multiple comparisons between ancestral phenotypic traits and the geometric mean of the number of adults and larvae evolving in corn per 10-generation transfer bin, computed across lines. Only statistically significant Spearman's rho correlation coefficients after correction are shown.

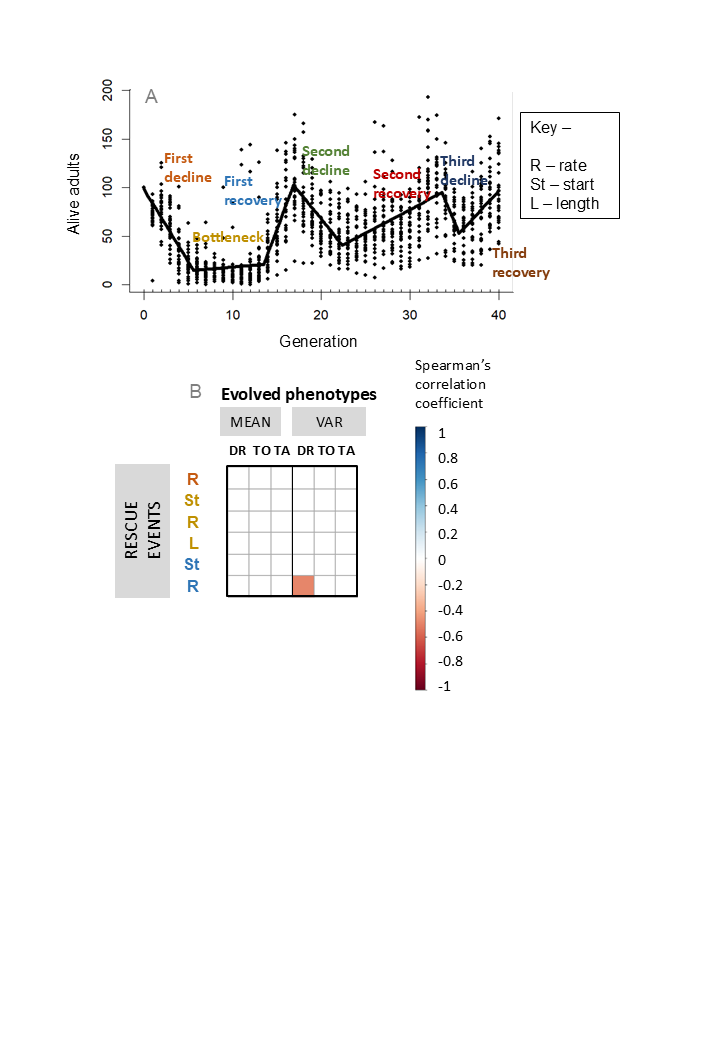

**Figure S8. Evolved phenotypic correlations (means and variances) with demographic rescue parameters, spread 50 generation apart.** (A) Rescue decline, bottleneck and recovery (rates, starts and lengths) in the context of other demographic events (shown for reference only) and (B) Matrices showing significant correlations corrected for multiple comparisons (blue – positive, red - negative) between the individual demographic events with evolved phenotypic traits, computed across lines. Only statistically significant Spearman's rho correlation coefficients after correction for multiple comparisons are shown. ‘Rates’ are extracted slopes from models fit individually to each line, ‘starts’ indicate the fitted breakpoint where one event ends and the other begins, and the bottleneck length. DR – development rate, TO – total offspring, TA – total adult offspring.

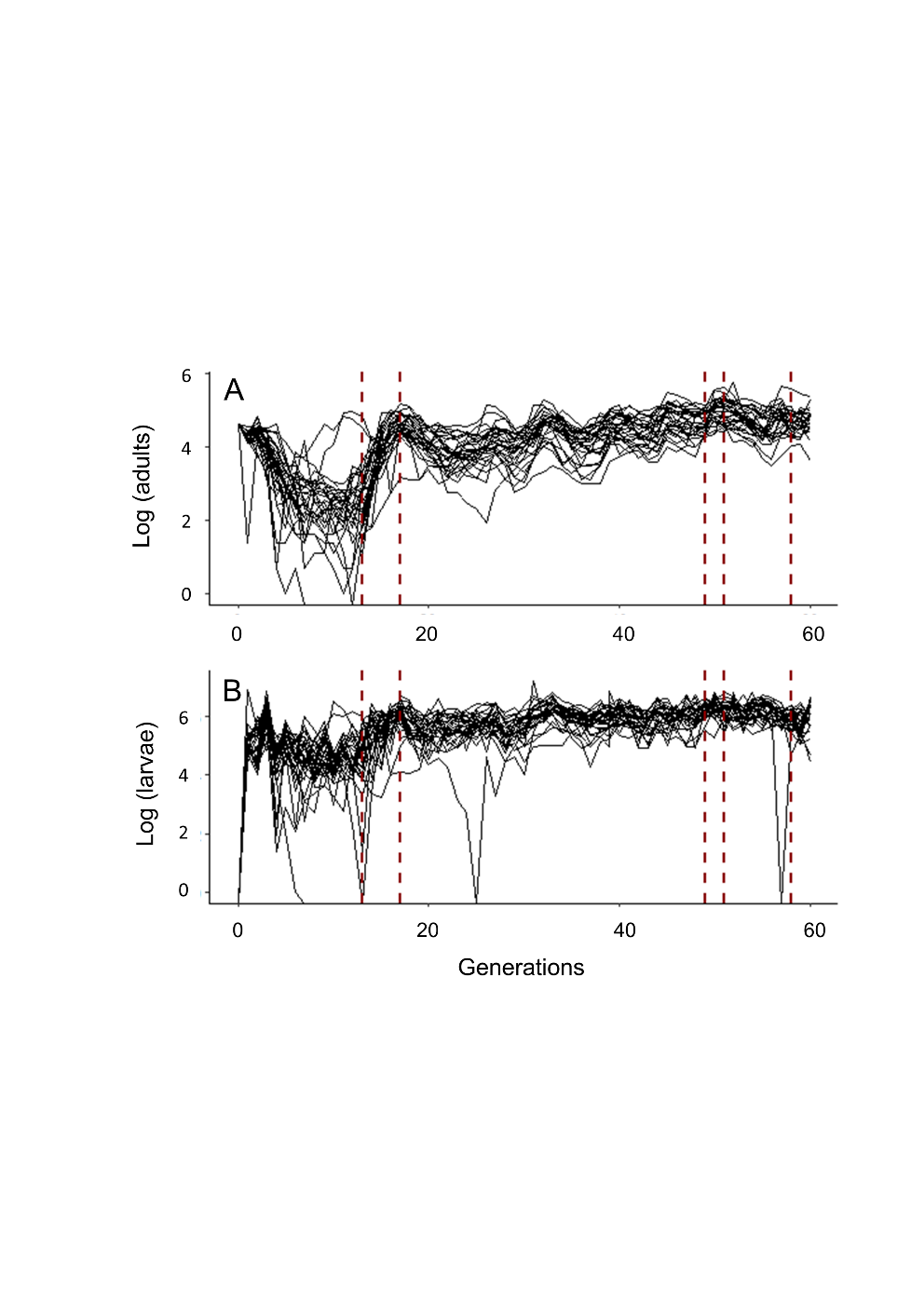

**Figure S9.** Time series of the (A) adult and (B) larval census in all the 30 selection lines (y-axis) over time (x-axis) overlaid with brown dashed lines indicating mild changes in the texture of the corn flour used to impose selection.

**SUPPLEMENTARY TABLES**

**Table S1.** Statistical summaries from tests for the effect of source population identity (“Pop”) on measured phenotypic traits. Model: phenotypic trait ~ source population.

| **Trait** | **Test** | **Factor** | **df** | **Test statistic** | **Proportion variance explained** | **p-value** |
| --- | --- | --- | --- | --- | --- | --- |
| **Longevity** | Cox PH | Pop | 9 | 32.26 | 0.05 (eta-squared) | 0.00018 |
| **Starvation resistance** | ANOVA | Pop | 8 | 9.618 | 0.08 (eta-squared) | 7.30e^-13^ |
|  |  | Residuals | 930 |  | | |
| **Larval cannibalism** | ANOVA | Pop | 9 | 2.934 | 0.1 (eta-squared) | 0.0025 |
|  |  | Residuals | 239 |  | | |
| **Development rate** | Binomial GLM | Pop | 9 | 89.43 (χ²) | 0.25 (Nagelkerke's R²) | 2.123e^-15^ |
| **Total offspring** | ANOVA | Pop | 9 | 7.775 | 0.15 (eta-squared) | 1.50e^-10^ |
|  |  | Residuals | 386 |  | | |
| **Total adult offspring** | Kruskal-Wallis | Pop | 9 | 16.13 (K-W H) | 0.02 (epsilon-squared) | 0.064 |

**Table S2.** Wilcoxon signed-rank exact test for change in average population size of each evolved line (for adults and larvae) between consecutive transfer bins of 10 generations each.

| **Stage** | **Comparison (generation-bins)** | **Sample estimate** | **95% confidence interval** | **p-value** |
| --- | --- | --- | --- | --- |
| Adults | 1 and 2 | -16.21377 | -20.74631 to -11.81912 | 1.028e^-5^ |
|  | 2 and 3 | -13.1047 | -21.013194 to -6.896064 | 6.048e^-5^ |
|  | 3 and 4 | -24.71332 | -30.18709 to -20.00566 | 7.451e^-9^ |
|  | 4 and 5 | -24.08426 | -30.89653 to -17.48478 | 5.215e^-8^ |
|  | 5 and 6 | -9.298974 | -13.887385 to -3.676717 | 0.001527 |
| Larvae | 1 and 2 | -88.68599 | -114.32461 to -62.82348 | 4.698e^-6^ |
|  | 2 and 3 | -68.7069 | -94.65099 to -40.61386 | 2.427e^-5^ |
|  | 3 and 4 | -117.8324 | -142.42668 to -89.22792 | 7.451e^-9^ |
|  | 4 and 5 | -6.323089 | -29.52630 to 15.52147 | 0.6239 |
|  | 5 and 6 | -25.61896 | -54.010075 to 6.514396 | 0.09633 |

**Table S3.** Wilcoxon signed-rank exact test for change in average coefficient of variation of each evolved line (for adults and larvae separately) between consecutive transfer bins.

| **Stage** | **Comparison (generation-bins)** | **Sample estimate** | **95% confidence interval** | **p-value** |
| --- | --- | --- | --- | --- |
| Adults | 1 and 2 | 15.66233 | 7.210736 - 23.207788 | 0.0007359 |
|  | 2 and 3 | 44.6446 | 36.43972 - 51.18989 | 2.608e^-8^ |
|  | 3 and 4 | -1.866084 | -7.293023 - 3.755448 | 0.4946 |
|  | 4 and 5 | -1.286663 | -5.592150 - 3.286994 | 0.5504 |
|  | 5 and 6 | 3.8951 | -1.131664 - 8.249127 | 0.1155 |
| Larvae | 1 and 2 | 19.95939 | 8.88337 - 32.39237 | 0.002542 |
|  | 2 and 3 | 27.55823 | 22.23024 - 33.28032 | 3.725e^-9^ |
|  | 3 and 4 | 1.662173 | -3.710282 - 7.117880 | 0.4946 |
|  | 4 and 5 | -2.245935 | -6.469355 - 2.772131 | 0.2941 |
|  | 5 and 6 | 0.6816072 | -4.877626 - 5.703017 | 0.848 |

**Table S4.** ANOVA summary statistics for average population size of each line as a function of source population identity in each transfer bin.

| **Stage** | **Generation-bin** | **SS** | **df** | **MS** | **F** | **p-value** |
| --- | --- | --- | --- | --- | --- | --- |
| Adults | 1 | 2017 | 9 | 224.14 | 2.424 | 0.0476 |
|  | 2 | 5113 | 9 | 568.2 | 3.202 | 0.0157 |
|  | 3 | 6271 | 9 | 696.8 | 2.63 | 0.0364 |
|  | 4 | 9129 | 9 | 1014.3 | 2.532 | 0.0423 |
|  | 5 | 11052 | 9 | 1227.9 | 1.593 | 0.188 |
|  | 6 | 8795 | 9 | 977.2 | 1.301 | 0.299 |
| Larvae | 1 | 12450 | 9 | 1383 | 0.512 | 0.849 |
|  | 2 | 107305 | 9 | 11923 | 4.945 | 0.00165 |
|  | 3 | 155307 | 9 | 17256 | 3.176 | 0.0163 |
|  | 4 | 134074 | 9 | 14897 | 2.695 | 0.033 |
|  | 5 | 107629 | 9 | 11959 | 2.226 | 0.068 |
|  | 6 | 192953 | 9 | 21439 | 2.85 | 0.0262 |

**Table S5.** ANOVA summary statistics for effect of source population on CV for each evolved line, tested separately for data in each transfer bin.

| **Stage** | **Generation-bin** | **SS** | **df** | **MS** | **F** | **p-value** |
| --- | --- | --- | --- | --- | --- | --- |
| Adults | 1 | 5137 | 9 | 570.8 | 1.459 | 0.23 |
|  | 2 | 2930 | 9 | 325.5 | 1.986 | 0.0996 |
|  | 3 | 1586 | 9 | 176.2 | 1.649 | 0.171 |
|  | 4 | 1090 | 9 | 121.17 | 1.506 | 0.216 |
|  | 5 | 675.2 | 9 | 75.02 | 0.67 | 0.726 |
|  | 6 | 871 | 9 | 96.78 | 1.809 | 0.132 |
| Larvae | 1 | 7229 | 9 | 803.2 | 1.085 | 0.415 |
|  | 2 | 3092 | 9 | 343.5 | 2.178 | 0.0734 |
|  | 3 | 2538 | 9 | 282.0 | 1.676 | 0.164 |
|  | 4 | 1881 | 9 | 209.0 | 1.895 | 0.115 |
|  | 5 | 1461 | 9 | 162.36 | 2.217 | 0.069 |
|  | 6 | 1236 | 9 | 137.36 | 1.588 | 0.189 |

**Table S6.** Spearman rank correlation statistics for differences in the average number of adults in consecutive transfer bins.

| **Stage** | **Comparison (generation-bins)** | **S** | **rho** | **p-value** |
| --- | --- | --- | --- | --- |
| Adults | 1 and 2 | 2436 | 0.4 | 0.03242 |
|  | 2 and 3 | 1026 | 0.7472906 | 6.593e^-06^ |
|  | 3 and 4 | 746 | 0.8162562 | 1.292e^-06^ |
|  | 4 and 5 | 918 | 0.7738916 | 2.677e^-06^ |
|  | 5 and 6 | 604 | 0.8512315 | 9.534e^-07^ |

**Table S7.** ANOVA summary statistics for the full model analysis for each phenotypic trait: phenotypic trait ~ source population x development resource.

| **Trait** | **Factor** | **SS** | **df** | **MS** | **F** | **Eta-squared** | **p-value** |
| --- | --- | --- | --- | --- | --- | --- | --- |
| Starvation resistance | Pop | 205 | 9 | 22.8 | 10.30 | 0.04 | 1.10e^-15^ |
|  | Dev resource | 620 | 1 | 620.1 | 279.95 | 0.12 | < 2e^-16^ |
|  | Interaction | 120 | 8 | 15.0 | 6.76 | 0.03 | 8.79e^-09^ |
| Larval cannibalism | Pop | 24.8 | 9 | 2.760 | 3.613 | 0.07 | 0.000233 |
|  | Dev resource | 5.4 | 1 | 5.355 | 7.009 | 0.02 | 0.008406 |
|  | Interaction | 9.9 | 9 | 1.097 | 1.437 | 0.03 | 0.169903 |
| Development rate | Pop | 6.42 | 9 | 0.714 | 10.71 | 0.14 | 1.72e^-15^ |
|  | Dev resource | 10.23 | 1 | 10.233 | 153.54 | 0.2 | < 2e^-16^ |
|  | Interaction | 7.94 | 9 | 0.883 | 13.25 | 0.17 | < 2e^-16^ |
| Total offspring | Pop | 780 | 9 | 86.63 | 7.912 | 0.09 | 3.37e^-11^ |
|  | Dev resource | 213 | 1 | 213.27 | 19.478 | 0.03 | 1.17e^-05^ |
|  | Interaction | 566 | 9 | 62.93 | 5.747 | 0.07 | 9.90e^-08^ |
| Total adult offspring | Pop | 53.8 | 9 | 5.97 | 12.98 | 0.14 | <2e^-16^ |
|  | Dev resource | 33.3 | 1 | 33.31 | 72.40 | 0.09 | <2e^-16^ |
|  | Interaction | 53.0 | 9 | 5.89 | 12.81 | 0.14 | <2e^-16^ |

**Table S8.** Spearman rank correlation statistics for differences in the fitness of ancestral populations that developed in either wheat (mimicking the founders) or corn (mimicking their generation 1 offspring) in the selection lines.

| **Trait** | **S** | **Spearman’s rho** | **p-value** |
| --- | --- | --- | --- |
| Starvation resistance | 142.68 | -0.1889 | 0.6263 |
| Larval cannibalism | 125.15 | 0.2415 | 0.5014 |
| Development rate | 149.74 | 0.0924 | 0.7995 |
| Total offspring | 160.61 | 0.0266 | 0.9418 |
| Total adult offspring | NA – too many zero values to compute | | |

**Table S9.** Summary statistics from a generalized linear model testing the difference in trait values between the ancestral populations (developed either in wheat – mimicking founders – or corn – mimicking their offspring) and each of the three evolved lines per source population.

| Trait | Population | Developed in | Evolved line 1 | | Evolved line 2 | | Evolved line 3 | |
| --- | --- | --- | --- | --- | --- | --- | --- | --- |
|  |  |  | Effect size | p-value | Effect size | p-value | Effect size | p-value |
| Development rate | Pop 1 | Wheat | 4.926717 | 1.03e^-12^ | 3.849282 | 5.25e^-10^ | 2.665726 | 3.99e^-5^ |
|  |  | Corn | 2.699682 | 8.08e^-12^ | 1.622248 | 7.12e^-11^ | 0.438691 | 0.163325 |
|  | Pop 2 | Wheat | 3.312803 | 5.81e^-37^ | Extinct | | 3.451848 | 2.01e^-22^ |
|  |  | Corn | 2.390996 | 8.91e^-35^ |  |  | 2.530041 | 2.51e^-16^ |
|  | Pop 6 | Wheat | 3.769019 | 2.91e^-43^ | 2.461311 | 3.08e^-26^ | 3.759105 | 3.06e^-60^ |
|  |  | Corn | 2.390268 | 1.34e^-27^ | 1.08256 | 6.47e^-11^ | 2.380354 | 6.87e^-49^ |
|  | Pop 7 | Wheat | 0.245235 | 0.36182 | 0.345908 | 0.174063 | 0.158224 | 0.690798 |
|  |  | Corn | 0.560996 | 0.003373 | 0.661669 | 0.000104 | 0.473985 | 0.175701 |
|  | Pop 11 | Wheat | 0.753118 | 0.021203 | 3.780802 | 9.47e^-41^ | 2.409988 | 2.87e^-22^ |
|  |  | Corn | 0.174516 | 0.540602 | 3.2022 | 8.54e^-43^ | 1.831387 | 6.16e^-22^ |
|  | Pop 12 | Wheat | 1.960703 | 9.29e^-13^ | 4.035504 | 3.77e^-21^ | 1.62885 | 1.53e^-7^ |
|  |  | Corn | 1.880555 | 1.82e^-13^ | 3.955356 | 1.74e^-21^ | 1.548702 | 1.31e^-7^ |
|  | Pop 13 | Wheat | 1.68279 | 3.01e^-13^ | 1.642281 | 3.32e^-13^ | 1.720892 | 1.12e^-13^ |
|  |  | Corn | 0.311352 | 0.048174 | 0.270843 | 0.07089 | 0.349455 | 0.028044 |
|  | Pop 18 | Wheat | 4.754206 | 7.35e^-13^ | 2.65926 | 8.94e^-10^ | 2.648789 | 2.27e^-10^ |
|  |  | Corn | 3.37588 | 2.09e^-9^ | 1.280934 | 6.92e^-7^ | 1.270463 | 3.10e^-8^ |
|  | Pop 20 | Wheat | -0.45611 | 0.365205 | 1.757376 | 1.20e^-10^ | 1.411934 | 1.09e^-7^ |
|  |  | Corn | -0.03323 | 0.944503 | 2.180259 | 4.47e^-23^ | 1.834817 | 4.30e^-18^ |
|  | Pop 24 | Wheat | 1.978671 | 2.93e^-8^ | 3.171975 | 1.21e^-18^ | 3.982905 | 1.55e^-30^ |
|  |  | Corn | -0.08645 | 0.671174 | 1.106851 | 1.18e^-7^ | 1.917781 | 4.59e^-25^ |
| Total offspring | Pop 1 | Wheat | 3.297727 | 0.000432 | 3.875 | 5.00e^-7^ | 3.297727 | 0.000432 |
|  |  | Corn | 2.243742 | 0.011172 | 2.821014 | 6.06e^-5^ | 2.243742 | 0.011172 |
|  | Pop 2 | Wheat | 5.12803 | 5.01e^-9^ | Extinct | | 4.408333 | 0.000203 |
|  |  | Corn | 4.519247 | 6.85e^-9^ |  |  | 3.79955 | 0.000677 |
|  | Pop 6 | Wheat | 3.041356 | 0.002286 | 1.728632 | 0.069242 | 5.700855 | 2.49e^-10^ |
|  |  | Corn | 2.664946 | 0.002589 | 1.352222 | 0.104966 | 5.324444 | 5.21e^-12^ |
|  | Pop 7 | Wheat | 5.939394 | 1.87e^-13^ | 8.636364 | 1.21e^-25^ | 3.701299 | 0.003315 |
|  |  | Corn | 4.484848 | 5.33e^-10^ | 7.181818 | 8.82e^-23^ | 2.246753 | 0.064196 |
|  | Pop 11 | Wheat | 4.525 | 0.000139 | 2.434091 | 0.000272 | 3.637069 | 1.53e^-6^ |
|  |  | Corn | 4.152778 | 0.000272 | 2.061869 | 0.000424 | 3.264847 | 1.69e^-6^ |
|  | Pop 12 | Wheat | 3.375375 | 0.098944 | 3.81982 | 0.062239 | -1.59685 | 0.38073 |
|  |  | Corn | 4.86072 | 0.013149 | 5.305164 | 0.006922 | -0.1115 | 0.947983 |
|  | Pop 13 | Wheat | 5.111888 | 3.88e^-12^ | 3.07089 | 1.89e^-6^ | 2.113782 | 0.001046 |
|  |  | Corn | 4.101702 | 2.37e^-10^ | 2.060704 | 0.000169 | 1.103596 | 0.045556 |
|  | Pop 18 | Wheat | 2.7 | 0.035542 | 3.9 | 3.71e^-6^ | 4.890476 | 2.59e^-10^ |
|  |  | Corn | 3.1125 | 0.012903 | 4.3125 | 5.55e^-8^ | 5.302976 | 1.90e^-13^ |
|  | Pop 20 | Wheat | 4 | 0.007281 | 4.515385 | 3.69e^-8^ | 6.490909 | 4.10e^-13^ |
|  |  | Corn | 3.1 | 0.031793 | 3.615385 | 7.50e^-7^ | 5.590909 | 4.52e^-12^ |
|  | Pop 24 | Wheat | 6.109023 | 6.44e^-7^ | 2.616959 | 0.016391 | 5.125506 | 3.08e^-7^ |
|  |  | Corn | 5.214286 | 4.76e^-6^ | 1.722222 | 0.084991 | 4.230769 | 2.26e^-6^ |
| Total adult offspring | Pop 1 | Wheat | 2.090909 | 2.11e^-8^ | 0.9 | 0.001842 | 1 | 0.005223 |
|  |  | Corn | 2.061924 | 7.74e^-9^ | 0.871014 | 0.001171 | 0.971014 | 0.004475 |
|  | Pop 2 | Wheat | 2.919697 | 1.26e^-19^ | Extinct | | 2.033333 | 6.51e^-7^ |
|  |  | Corn | 2.658886 | 2.82e^-20^ |  |  | 1.772523 | 3.96e^-6^ |
|  | Pop 6 | Wheat | 4.271299 | 2.01e^-11^ | 1.226496 | 0.03602 | 5.541311 | 2.92e^-21^ |
|  |  | Corn | 3.549247 | 2.70e^-10^ | 0.504444 | 0.323803 | 4.819259 | 4.78e^-22^ |
|  | Pop 7 | Wheat | 1.108586 | 5.07e^-8^ | 0.765152 | 4.77e^-5^ | 0.600649 | 0.065816 |
|  |  | Corn | 1.131313 | 1.73e^-9^ | 0.787879 | 4.13e^-6^ | 0.623377 | 0.049214 |
|  | Pop 11 | Wheat | 0.75 | 0.239778 | 3.386364 | 1.47e^-17^ | 1.896552 | 4.29e^-6^ |
|  |  | Corn | 0.652778 | 0.287487 | 3.289141 | 1.70e^-20^ | 1.79933 | 1.42e^-6^ |
|  | Pop 12 | Wheat | 1.057057 | 2.73e^-6^ | 1.501502 | 1.40e^-10^ | 0.529279 | 0.006881 |
|  |  | Corn | 1.068858 | 7.17e^-7^ | 1.513302 | 1.62e^-11^ | 0.54108 | 0.003353 |
|  | Pop 13 | Wheat | 1.088578 | 0.001493 | 1.493213 | 1.76e^-6^ | 0.908654 | 0.003574 |
|  |  | Corn | 0.598589 | 0.04748 | 1.003223 | 0.000159 | 0.418664 | 0.11706 |
|  | Pop 18 | Wheat | 3.2 | 4.60e^-22^ | 0.333333 | 0.066587 | 0.285714 | 0.077179 |
|  |  | Corn | 3.1875 | 7.82e^-23^ | 0.320833 | 0.057537 | 0.273214 | 0.063395 |
|  | Pop 20 | Wheat | -0.05 | 0.89333 | 0.988462 | 1.45e^-6^ | 1.404545 | 2.47e^-10^ |
|  |  | Corn | -0.02564 | 0.943631 | 1.012821 | 5.40e^-8^ | 1.428904 | 2.89e^-12^ |
|  | Pop 24 | Wheat | 0.214286 | 0.598101 | 0.722222 | 0.05349 | 2.884615 | 2.82e^-15^ |
|  |  | Corn | -0.24405 | 0.520582 | 0.263889 | 0.441432 | 2.426282 | 8.03e^-14^ |

**Software and Packages Used**

Auguie, B. and Antonov, A. (2017) gridExtra: Miscellaneous Functions for “Grid” Graphics. R Package Version 2.3. <https://CRAN.R-project.org/package=gridExtra>

Garnier, Simon, Ross, Noam, Rudis, Robert, Camargo, Pedro A, Sciaini, Marco, Scherer, Cédric (2024). viridis(Lite) - Colorblind-Friendly Color Maps for R. doi:10.5281/zenodo.4679423, viridis package version 0.6.5, <https://sjmgarnier.github.io/viridis/>.

Hvitfeldt E (2021). paletteer: Comprehensive Collection of Color Palettes. R package version 1.3.0, <https://github.com/EmilHvitfeldt/paletteer>.

Kassambara A (2023). ggpubr: 'ggplot2' Based Publication Ready Plots. R package version 0.6.0, <https://rpkgs.datanovia.com/ggpubr/>.

Lüdecke D (2024). sjstats: Statistical Functions for Regression Models (Version 0.19.0). doi:10.5281/zenodo.1284472, <https://CRAN.R-project.org/package=sjstats>.

Muggeo VM (2008). “segmented: an R Package to Fit Regression Models with Broken-Line Relationships.” R News, 8(1), 20–25. <https://cran.r-project.org/doc/Rnews/>.

Neuwirth, E. (2014) RColorBrewer: ColorBrewer Palettes. R Package Version 1.1-3.

<https://CRAN.R-project.org/package=RColorBrewer>

Oksanen J, Simpson G, Blanchet F, Kindt R, Legendre P, Minchin P, O'Hara R, Solymos P, Stevens M, Szoecs E, Wagner H, Barbour M, Bedward M, Bolker B, Borcard D, Borman T, Carvalho G, Chirico M, De Caceres M, Durand S, Evangelista H, FitzJohn R, Friendly M, Furneaux B, Hannigan G, Hill M, Lahti L, McGlinn D, Ouellette M, Ribeiro Cunha E, Smith T, Stier A, Ter Braak C, Weedon J (2025). vegan: Community Ecology Package. R package version 2.7-0, https://github.com/vegandevs/vegan, <https://vegandevs.github.io/vegan/>.

R Core Team (2021). R: A language and environment for statistical computing. R Foundation for Statistical Computing, Vienna, Austria. URL <https://www.R-project.org/>.

Robinson D, Hayes A, Couch S (2024). broom: Convert Statistical Objects into Tidy Tibbles. R package version 1.0.7, https://github.com/tidymodels/broom, <https://broom.tidymodels.org/>.

RStudio Team (2020). RStudio: Integrated Development for R. RStudio, PBC, Boston, MA URL <http://www.rstudio.com/>.

Rudis B (2025). hrbrthemes: Additional Themes, Theme Components and Utilities for 'ggplot2'. R package version 0.90, <https://github.com/hrbrmstr/hrbrthemes>.

Therneau T (2024). A Package for Survival Analysis in R. R package version 3.8-3, <https://CRAN.R-project.org/package=survival>.

Terry M. Therneau, Patricia M. Grambsch (2000). Modeling Survival Data: Extending the Cox Model. Springer, New York. ISBN 0-387-98784-3.

Venables WN, Ripley BD (2002). Modern Applied Statistics with S, Fourth edition. Springer, New York. ISBN 0-387-95457-0, <https://www.stats.ox.ac.uk/pub/MASS4/>.

Wickham H (2016). ggplot2: Elegant Graphics for Data Analysis. Springer-Verlag New York. ISBN 978-3-319-24277-4, <https://ggplot2.tidyverse.org>.

Wickham H, François R, Henry L, Müller K, Vaughan D (2023). dplyr: A Grammar of Data Manipulation. R package version 1.1.4, https://github.com/tidyverse/dplyr, <https://dplyr.tidyverse.org>.

Wickham H (2023). forcats: Tools for Working with Categorical Variables (Factors). R package version 1.0.0, https://github.com/tidyverse/forcats, <https://forcats.tidyverse.org/>.

Wickham H, Vaughan D, Girlich M (2024). tidyr: Tidy Messy Data. R package version 1.3.1, https://github.com/tidyverse/tidyr, <https://tidyr.tidyverse.org>.

Wei T, Simko V (2024). R package 'corrplot': Visualization of a Correlation Matrix. (Version 0.95), <https://github.com/taiyun/corrplot>.
